# Genomic insights into photosymbiotic evolution in *Tridacna squamosa*

**DOI:** 10.1101/2024.02.04.577604

**Authors:** Yang Zhang, Fan Mao, Yuanning Li, Nai-Kei Wong, Yongbo Bao, He Dai, Jin Sun, Wenjie Yi, Shu Xiao, Zhiming Xiang, Jun Li, Yuehuan Zhang, Xiaomin Xia, Lvping Zhang, Huawei Zhou, Ziniu Yu

## Abstract

Photosymbiosis is fundamental driving force for ecological success of benthic coral reef ecosystems, and contributing to their biodiversity and resilience. As a benchmark organism indicative of reef health, the fluted giant clam (*Tridacna squamosa*) forms an exemplary photosymbiotic relationship with the symbiont Symbiodiniaceae dinoflagellates, whose initiation and maturation require finely coordinated interactions. However, much of the origin and dynamics of this reciprocal interplay remains unclarified. Here, we report the first complete whole genome of *T. squamosa*, in conjunction with integrated multi-omics data, to illuminate the key evolutionary innovations and molecular events supporting the establishment and maintenance of photosymbiotic lifestyle in the giant clam. Programmed regulation of symbiont recognition, host immune system and GPCRs signaling activation co-contributed to dinoflagellates acquisition in *T. squamosa* larvae. Adaptive metabolic remodeling in the host siphonal mantle, a photosymbiotic niche, is critical to maintain the robustness of phtosymbiosis. *T. squamosa* has expanded light sensing gene family and evolved sophisticated signaling pathways to protect against UV photo-damage. Evidence also supports significant contribution of positive selection to host DNA-repair. Overall, our study here offers fresh mechanistic insights into the parallel evolution and molecular machinery of photosymbiosis in the giant clam-dinoflagellates duet, with implications for devising solutions to sustainable conservation.

## Introduction

Photosymbiosis is an extraordinary phenomenon in which interspecies communication and biosynthetic interplay co-occurs in exquisite balance. In marine habitats, photosynthetic microorganisms (symbionts) such as Symbiodiniaceae dinoflagellates dwelling within heterotrophic host organisms play ecologically pivotal roles in shaping biodiversity and evolution of benthic coral reef ecosystems ^1^. Indeed, coral reefs ecosystems are home to the world’s maximal marine biodiversity (up to 25%) ^2^, but their own survival remains threatened by climate change and anthropogenic encroachment ^3, 4, 5^. Among coral reef flora and fauna, giant clams (*Tridacna* spp.), the largest and prominent tropically distributed bivalves, have evolved unique symbiotic relations with photosynthetic dinoflagellates to thrive in oligotrophic (or nutrient-deficient) marine environments, which spatially overlap with Indo-Pacific coral reefs ^6, 7^. In terms of ecological relevance, giant clams are essential to the integrity of healthy coral reefs, with such underappreciated roles such as reef builders and shapers via furnishing a calcium carbonate framework, and contributors of coral reef resilience and diversity by providing shelter or food for other commensal and ectoparasitic organisms^7^.

Remarkably, giant clams also boost anti-biofouling capacities of marine habitats as they physically form assemblages with corals ^8^. Unlike the case of intracellular symbiosis in cnidarian corals ^9^, giant clams possess specialized symbiotic tissues known as the “zooxanthellal tubular system” to accommodate dinoflagellates in their siphonal mantle ^10^, where the host provides shelter for symbiont dinoflagellates against predation, while the symbiont produces and transfers the majority of its photosynthesis substrates to the host in reciprocity ^11^. Such evolutionary and ecological success of photosymbiotic species, like corals or giant clams, in nutrient-limited tropical waters, have been frequently referred to as a “Darwin’s paradox” ^12^. Despite extensive efforts in investigating the elusive nature of cnidarian-dinoflagellate symbiosis, there remain huge gaps in our knowledge with respect to the origin and mechanistic aspects of photosymbiosis ^9, 13^. Intuitively, the giant clam-dinoflagellates holobiont provides an excellent model for elucidating parallel evolution and diversity of photosymbiosis in metazoans.

Climate change and accompanying decay of marine ecosystems have made photosymbiosis research an urgent priority for better understanding the loss of marine biodiversity. Despite being designated on the IUCN red list of threated species and CITES (Convention on International Trade in Endangered Species) Appendix II, global populations of giant clams remain in constant peril, with even risks of extinction in many native regions of their distributions. Major causes for this crisis include excessive commercial exploitation for highly prized clam shells, the organism’s vulnerability to climate perturbations, and falling fecundity in the hermaphroditic reproductive cycles ^14, 15^. To this date, 12 of the existing giant clam species have been studied, including 10 of Tridacna genus and 2 of the Hippopus genus ^16^. Among these, *Tridacna squamosa* (fluted giant clam), is physically one of the largest species native to tropical coral reefs of the Indo-Pacific ^17^. Recently, efforts on molecular underpinnings of the fluted giant clam-dinoflagellate association have revealed the involvement of inorganic carbon or nitrogen assimilation, and light-dependent enhancement of various transporters or enzymes ^13^. However, these studies are discrete and limited in scope with several known genes. Herein, we report comprehensive analyses on the first complete whole genome of the giant clam, in conjunction with insights from integrated multi-omics, to reveal key evolutionary innovations and mechanistic details on the establishment and maintenance of photosymbiosis, light sensing and light protection of the host.

## Results and discussion

### Section 1: Genome landscape of the giant clam

1. *T. squamosa* is one of representative native giant clams with distribution among the tropical Indo-Pacific coral reefs (**Figure 1A**). By using PacBio RSII single-molecule real-time (SMRT) sequencing technology, 88% coverage of PacBio sequencing reads was sequenced and assembled into a 1.08 Gb genome (**Supplementary Table 1**), which is approximately equivalent to an estimated genome size of 1.09 Gb by k-mer analysis (**Figure S1**). Of this, 1.05 Gb of the genome was anchored onto 18 pseudo-chromosomes with a scaffold N50 of 59.43 Mb (**Figure 1B and Figure S2A; Supplementary Table 1 & 2**), being consistent with the results for karyotype analysis (**Figure S2B**). Quality evaluation of the initial genome assembly suggests that 99.14% and 98.93% of the Illumina reads from two samples were successfully mapped onto the assembled genome (**Supplementary Table 3**). Genome integrity was assessed to be 95.89%-99.14% of the sequencing reads mapping (**Supplementary Table 3**) and 94.97% of BUSCO completeness (**Supplementary Table 4**), which is superior in genome completeness to *Ruditapes philippinarum* (93.61%) ^18^ and *Sinonovacula constricta* (88.88%) ^19^ as analyzed comparatively in this study. For gene annotation, 24,023 protein-coding genes were predicted in the genome by integrating results from *ab initio* prediction, homology-based searches with reference genomes and RNA-seq (**Supplementary Table 5**).

**Figure 1.**
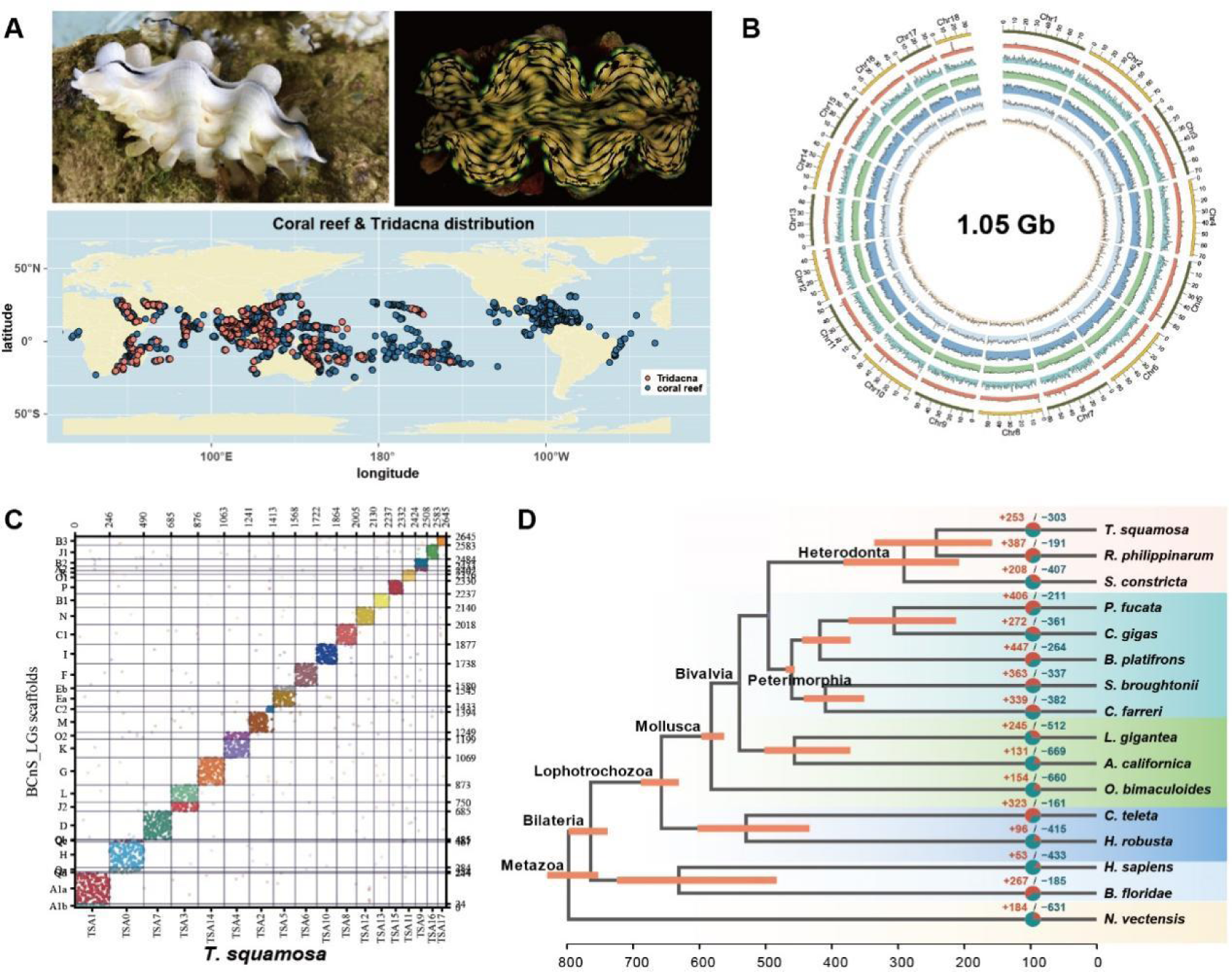
Geographical niches, genome landscape, and phylogenetic analysis of the *T. squamosa*. **A,** Global distribution of giant clam (red dots) and coral reef (blue dots). **B,** Circos plot highlights genome characteristics across 18 chromosomes in a megabase (Mb) scale. The sequencing depth, gene density, GC content percentage, LTR, LINE, and SINE are presented from outer to inner circles. **C,** Chromosome-based macrosynteny is shown in the form of Oxford dot plots with comparisons between the chromosomes of *T. squamosa* (*x* axis) and the previously reconstructed 29 BCnS-ALGs (*y* axis). **D,** A phylogenetic tree was constructed across T. squamosa and 15 other metazoan animals based on 423 orthologues using OrthoMCL with a Markov cluster algorithm. The positive and negative numbers adjacent to the taxon names are gene family numbers of expansion/contraction obtained from the CAFE analysis.

Comparative analysis further revealed that the genome of *T. squamosa* exhibits the smallest size among its heterodonata congeners, while concurrently possessing the longest introns (**Supplementary Table 6**). Moreover, predicted repetitive elements occupies a collective length of 628.34 Mb, accounting for 58.07% of the genome (**Supplementary Table 7**). Furthermore, distinct long terminal repeats (LTR) make up the largest portion of the repetitive sequences (**Figure S3**), and species specifically expanded in the *T. squamosa* genome when compared to closed relatives *R. philippinarum* and *S. constricta* by Kimura distance-based copy divergence analysis (**Figure S4**), implicating the critical role of retrotransposons in driving rapid genome evolution in this symbiotic bivalve species. Furthermore, to retrace the history time of two major expansion waves of LTR in *T. squamosa* genome, TEs were dated with RepeatedMasker and adjusted by using Jukes-Cantor formula ^20^, which revealed two major expansion waves of LTR in *T. squamosa* genome at 0.13 and 0.23 (**Figure S4**). Through comparison to the dS measure and estimation for synonymous substitutions per million years (**Figure S5**), it was determined that JK 0.13 and 0.23 correspond to 28 and 50 million years ago (MYA), respectively, whereby the first engagement in symbiotic association of the Tridacna giant clam-dinoflagellates pair was posited to occur during this Eocene-Oligocene transition period ^11^. The evidence here implies that transponson propagation may serve as the main driving force to shape genome innovations underpinning symbiosis in the giant clam. In addition, comparative analysis of chromosomal structures revealed a significant degree of macrosynteny conservation between *T. squamosa* and other species despite an evolutionary divergence exceeding 600 million years (**Figure 1C & Figure S7& Figure S6**), suggesting a remarkably conservative karyotype without significant change. As support to this, a complete molluscan Hox geneset was identified in *T. squamosa* genome, which is in agreement with existing evidence that molluscan ancestors preserved intact Hox clusters 21, 22.

To illuminate the phylogenetic relationship between *T. squamosa* and other species, OrthoMCL was used to identify the gene orthologous of 16 species, which generated 423 single-copy orthologous for further construction of phylogenetic tree (**Supplementary Table 8**). Phylogenetic analysis confirmed that the giant clam diverged from other heterodonata congeners at around 240.92 MYA (**Figure 1D**). Analysis of gene families based on CAFÉ ^23^ indicates that 253 orthologous gene families were significantly expanded, whereas 303 orthologous gene families were contracted in *T. squamosa* (**Figure 1D**). Specially, KEGG enrichment analysis of the gene family further shows that molecular signaling and phototransduction pathways were strictly enriched in gene expanded families, while several metabolic pathways were contracted in the host (**Figure S9, Supplementary Table 9 & 10**), strongly corroborating the idea that signaling pathways and metabolic activities have been fundamentally reshaped following the advent of photosymbiotic life in *T. squamosa*.

### Section 2: Symbiosis establishment

Establishment of obligate symbiotic association with dinoflagellates are mandatory and essential for survival of photosymbiotic reef animals. Thus, in each life cycle, the juvenile Tridacna giant clam needs to acquire the dinoflagellates afresh from the marine environment to reinitiate a symbiotic relationship at the early veliger stage, failing which, these giant clams will invariably perish for being unable to forge a productive association with the symbiont ^24, 25^. To dissect the molecular mechanisms underlying the onset of symbiotic establishment in the giant clam, dual time course experiments were performed (start from early veliger stage and persistence of 72 h). Specifically, a productive group contained symbiotic larvae for colonization with native Symbiodinium *spp.* dinoflagellates, while an unproductive group contained nonsymbiotic larvae fed with chrysophyceae (*Isochrysis galbana*) as a control (**Figure 2A**). Totally, 22 mRNA libraries from 6 time points were sequenced, followed by time-series gene expression analysis by using the maSigPro R package ^26^ (**Figure S10**). The results showed that a total of 20.42% of genome encoding genes during the symbiotic establishment were dynamically altered, including early 1,759 DGEs and late 1,260 DGEs in symbiotic larvae, and 1,280 early DGEs and 607 late DGEs in nonsymbiotic larvae, respectively (**Figure 2B & Figure S10**), suggesting that massive shifts in gene expression is phenotypically accompanied by lifestyle conversion from heterotrophs to autotrophs in giant clam larvae.

**Figure 2.**
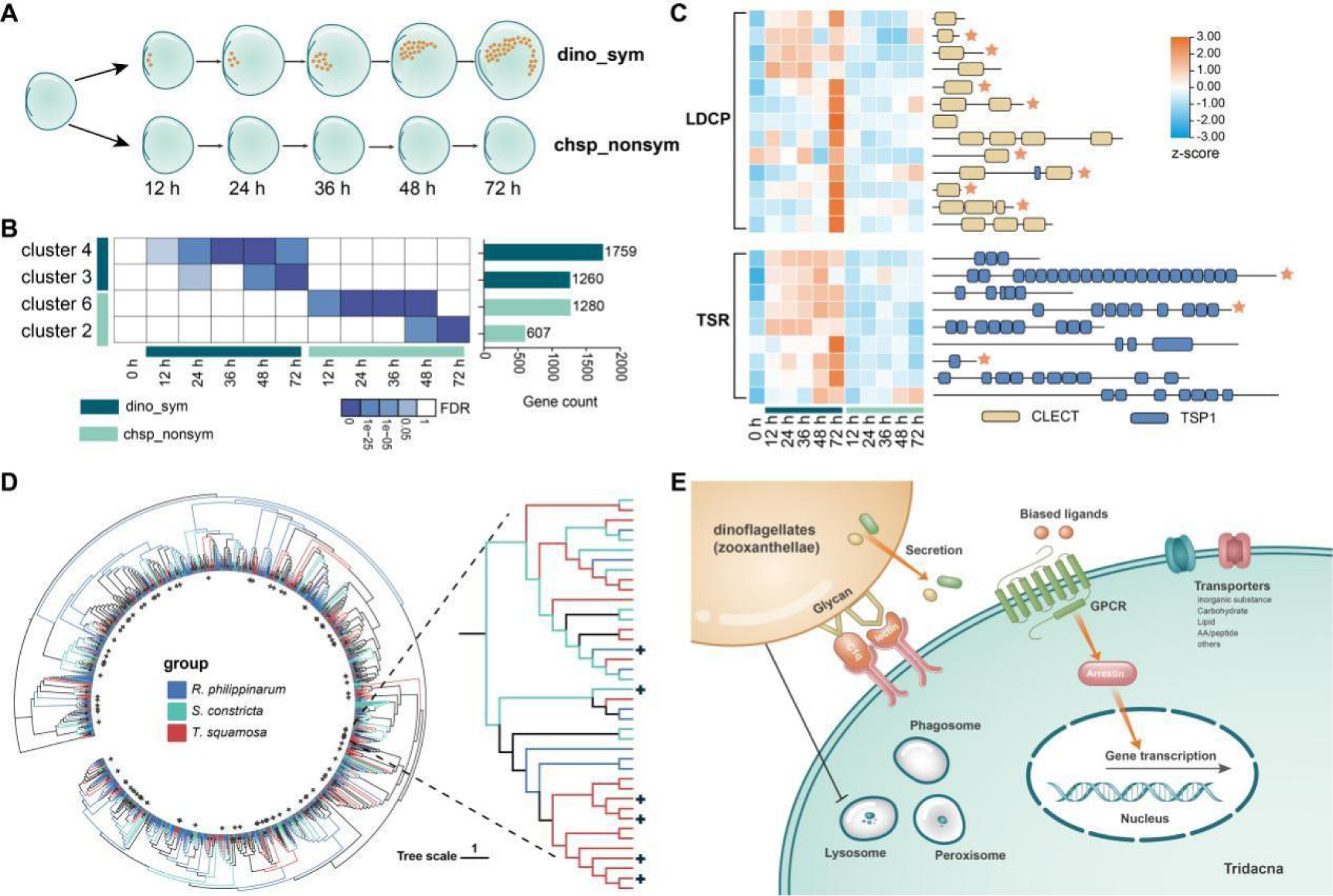
Molecular interactions involved in Tridacna-Symbiodiniaceae symbiosis establishment. **A,** Schematic overview of symbiotic and nonsymbiotic algae feeding duration. Dino_sym is short for *Symbiodinium spp.* dinoflagellates (zooxanthellae) feeding larvae, while chsp_nonsym is short for chrysophyceae feeding larvae. **B,** Deferentially expressed gene datasets analyzed by MaSigPro package. Gene numbers are indicated as histogram at right panel. **C,** Heatmap indicates the expression level of part of Lectin domaing containing proteins (LDCPs) and thrombospondin type 1 repeat (TSRs) involved in symbiosis establishment as pattern recognition receptors, along with their protein structure. Pentagram marks the proteins that contain the signal peptide. **D,** Phylogenetic tree of GPCRs encoded in the *T. squamosa* genome with other closed Heterodonta bivalve species constructed by maximum likelihood method. The “+” symbol marks the GPCR genes specifically expressed in *T. squamosa* during symbiotic dinoflagellates feeding. **E,** Schematic diagram of the molecular and pathways supposed to trigger Tridacna-Symbiodiniaceae symbiosis establishment

Generally, the first steps of success toward symbiotic establishment entail highly specific symbiont recognition, which is governed by microbe-associated molecular pattern (MAMP)-pattern recognition receptor (PRR) interactions ^9^. In cnidarians, a large body of evidence has demonstrated that lectins, Toll-like receptors (TLRs), scavenger receptors (SRs), and thrombospondin type 1 repeats (TSRs) participate in MAMP-PRR interactions during symbiosis establishment ^9, 27, 28^. Similarly, we observed that 162 of lectin domain containing proteins (LDCPs), 62 of TLRs, 51 of TSRs, and 32 of SRs were encoded in the *T. squamosa* genome (**Supplementary Table 11**), implying that diversity of recognition receptors may be functionally central to in giant clam-dinoflagellates association. It is worth noting that 13 of LDCPs and 9 of TSRs are activated during the early and late stages of colonization of dinoflagellates (**Figure 2C**), highlighting their essential roles in conferring host-symbiont specificity. Additionally, majority of LDCPs and TSRs were predicted to possesses a signal peptides domain (**Figure S11**), providing further evidence for secretion and binding of glycosaminoglycans on the surface of dinoflagellates *in hospite*. In contrast, TLRs and SRs, which have been considered to play a role in immune response ^29, 30^, were increased in transcripts in nonsymbiotic rather symbiotic *T. squamosa* larvae, (**Figure S12**), suggesting that selective recognition receptors activation may contribute the specific symbiotic algae *in hospite* colonization.

Following symbiont recognition, the cnidarian host is licensed to engulf dinoflagellates through intracellular phagocytosis and subsequently installs an endosymbiosis lifestyle, for which hypothetical immune suppression and arrest of phagosome maturation have been anticipated to be a prerequisite for the establishment of compatible symbioses ^31^. By comparison, Tridacna giant clam veliger larvae is capable of taking up and retaining dinoflagellates in the host digestive tract, and then migrating them into the primordial siphonal mantle 72 h after acquisition (**Figure 2A**). During this whole process, the host Tridacna giant clam did not phagocytize dinoflagellates symbiont and form extracellular association ^11^. It is thus intriguing to ask whether the previous assumption of “symbiosis derived immune suppression” could be applied in the giant clam. Surprisingly, KEGG enrichment analysis showed that several signaling pathways including “lysosome (ko04142, *p* = 1.64E-11)”, “phagosome (ko04145, *p* = 6.43E-5)” and “peroxisome (ko04146, *p* = 0.0001)” involved in immune response and phagosome maturation are downregulated during dinoflagellates colonization in symbiotic *T. squamosa* larvae, as compared with that in nonsymbiotic larvae (**Figure S13**), suggesting that selective suppression of intrinsic immune system could be a conserved feature of host facilitating mechanism underlying symbiont colonization in Cnidarians as well as in Tridacna.

Moreover, our results showed that active transcripts of gene are significantly enriched in 69 and 99 signaling pathways in early and late symbiotic *T. squamosa* larvae, respectively (**supplementary table 12 & 13**). Strikingly, the most enriched pathway “transporter” (*p=* 0.003 for early symbiosis and *p=* 0.0002 for late symbiosis) are persistently active during dinoflagellates colonization, supporting the idea that metabolite transfer from the symbiont to the host may be initiated from early stage upon dinoflagellates acquisition. Furthermore, a comprehensive analysis on tissue distribution patterns of transporters in adults revealed that a significant portion of these dominating expression transporters genes (including sodium-dependent glucose transporter, amino acid transporter, sodium-coupled monocarboxylate transporter, etc.) during dinoflagellates colonization persist in an active state in the siphinol mantle, which is a specific symbiotic tissue in adult giant clams (**Figure S14**). This strongly suggests that metabolic remodeling of the transporter toolkit is critical to global shifts from transitional to stable symbiosis. Consistent with this idea, specific signaling pathways were observed to be activated in early and late symbiotic *T. squamosa* larvae, respectively (**Figure S13**). For example, the transmembrane G protein-coupled receptors (GPCRs) are capable of transducing extracellular signals into pleiotropic effects by sensing and binding of various endogenous ligands, including hormones, neurotransmitters, and metabolites such as carbohydrates, lipids, amino acids, peptides, and proteins ^32^. Furthermore, phylogenetic analysis of 544 GPCRs encoded in the *T. squamosa* genome with other closed Heterodonta bivalve species revealed a high genetic diversity and slightly lineage-specific expansions (**Figure 2D**). Among them, 81 and 21 of G protein-coupled receptors (GPCRs) are significantly upregulated in in the early and late stages of dinoflagellates colonization, whereas only 16 and 2 GPCRs are compared with those in early and late nonsymbiotic larvea (**Figure S15A**). This suggests the possible presence of a dynamic and complex metabolite sensing machinery at work before stable symbiotic establishment. In particular, four members within one lineage-specifically expanded GPCR subfamily (GPCR126) were observed to be upregulated (**Figure 2D**), implying that specific substance or metabolite sensing may contribute to lineage specific symbiosis establishment, which warrants further investigation. Consistently, a set of *β*-arrestin, the core regulator of GPCR signaling, are activated in conjunction with GPCRs during dinoflagellates colonization (**Figure S15B**), highlighting the functional significance of GPCRs signaling in symbiotic establishment. Overall, multiple molecular events including molecular patterns recognition, host immune suppression, arrest of phagocytosis maturation, and dynamic transporter and GPCRs signaling remodeling seem to operate in a programmed manner to drive symbiotic establishment in *T. squamosa* larvae (**Figure 2E**).

### Section 3: Symbiosis robustness

Metabolic exchange is central to the evolutionary success of photosymbiotic species. It is also essential to maintaining hemostasis in the holobiont and even adaption to climate change ^33, 34^. In the cnidarian-dinoflagellate symbiosis, the algal symbiont could acquire dissolved inorganic nutrients like carbon, nitrogen and phosphorous etc. and respiration-derived metabolites from host ^9^. In reciprocity, dinoflagellate symbiont provides the products of carbon fixation and nitrogen assimilation, including carbohydrates, amino acids and lipids, to host ^35, 36^. Such patterns of reciprocal exchange have been envisaged to be conserved within the lineage of Tridacna, though the underlying mechanisms therein remain obscure ^13^.

To this end, comparative genomic analysis was first conducted to identify the transporters governing metabolic translocation, which revealed a complete framework of transporters responsible for life-sustaining metabolites including carbohydrates/lipids, amino acids, vitamins and phosphate/sulphur elements being present in the giant clam genome (**Figure 3A, Supplementary Table 14**). However, in contrast to other non-symbiotic metazoan genomes, no significant transporter gene expansion except for one gene family was observed in the giant clam genome (**Figure 3A)**, leading to the hypothesis that transcriptional plasticity rather than transporter gene expansion is leveraged to maintain the giant clam-dinoflagellate symbiotic relationship. To test this hypothesis, we conducted transcriptomic analyses on multiple tissues, including the symbiotic mantle and other non-symbiotic organs. WGCNA (weighted gene co-expression network analysis) results showed that specific gene clusters are highly associated with each tissue (**Figure S16**). Remarkably, 188 of genes were observed to be specifically expressed in the symbiotic mantle, where the category “transporters” is primarily enriched based on gene annotation analysis (**Figure S17 & 18, Supplementary Table 15**), thus supporting our conjecture that transcriptional activation of inherent transporters in the symbiotic tissue is a principal strategy for achieving photo-symbiosis success in the giant clam.

**Figure 3.**
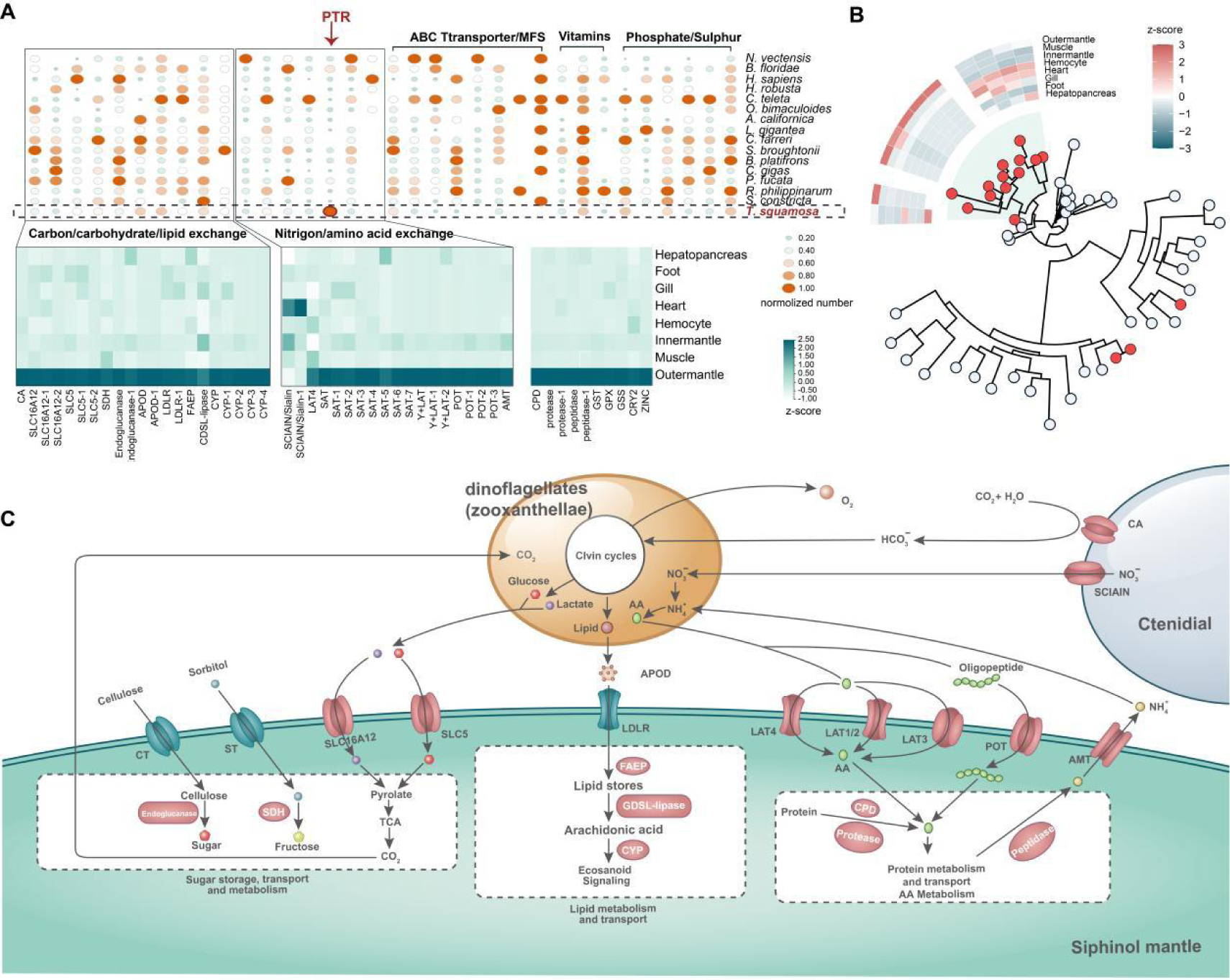
Substance transport system maintaining Tridacna symbiosis. **A,** Bubble heatmap illustrates the gene numbers of Pfam entries mostly associated with sunbstance exchange from *T. squamosa* and 15 other metazoan animals genome. For Pfam entries, corresponding ID and names are omitted, instead of the substance type they supposed to transport. PTR, proton-dependent oligopeptide transporter (POT). The heatmap below shows the expression level of corresponding transporter genes in different tissue, as *z*-score calculated from RPKM-values. **B,** Phylogenetic tree of POT gene family identified from *T. squamosa* and 15 other metazoan animals. Sequences from *T. squamosa* genome are displayed in red circles. The background in light-green color highlights the POT genes specifically expanded in *T. squamosa* genome. Heatmap of the expanded *Ts*POTs is also displayed as *z*-score calculated from RPKM-values. **C,** Schematic summary of substance exchange and events presumably occurring in symbiosis maintenance. All genes indicated are specifically expressed in symbiotic tissue, the siphonal mantle.

Carbohydrates including glucose and its catabolites are the primary carbon source of photosynthesis and are generally considered a major transferrable metabolite class to the host of coral or the giant clam ^35, 37^. In agreement with this, three members of SLC5 (solute carrier family 5) subfamily transporting glucose ^38^ and three members of SCL16 subfamily transporting lactate/pyruvate ^39^ were found to be predominantly expressed in the symbiotic mantle. Lipids could be further produced in the symbiont through utilization of inorganic carbon and carbohydrates in photosynthesis, which have recently been proven to be another main carrier of carbon translocation in the dynamic symbiotic relationship ^40, 41^. In this study, genomic and transcriptomic analyses revealed an active APOD-dependent LDLR lipid transport pathway in the symbiotic mantle (**Figure 3A**), implicating lipids as a key molecular class in carbon translocation supporting the giant clam-dinoflagellate association. Moreover, the genes involved in lipid storage and metabolism, such as GDSL-lipase and FAEP, were also observed to be specifically expressed in the symbiotic mantle, further strengthening our assumptions.

Apart from carbohydrates and lipids, multiple transporters responsible for nitrogen acquisition were also observed with specific expression patterns in the symbiotic mantle, including 9 of *L*-type amino acid transporter (LAT) and 4 of proton-coupled peptide transporters (POT/PTR family). Of note, the POTs (PF00854.20 PTR2) are the only expanded gene family with 16 POT genes encoded in *T. squamosa* genome in contrast with no more than 8 genes found in other bivalves (**Figure S19**), facilitating nutrient uptake by recognizing and transporting over 8,000 distinct dipeptide and tripeptide ligands ^42, 43^. Phylogenetic analysis further revealed that 13 of 16 POTs genes are clustered into one clade, supporting lineage-specific POTs gene expansion in the giant clam (**Figure 3B**). Generally, tandem duplication, segmental duplication, and whole-genome duplication are some of the primary driving forces for expanding gene families in specific species [31].Herein, we observed that 14 POTs are tandemly localized into two synteny blocks (**Figure S20**), suggesting that gene tandem duplication in combination with genomic fragment duplications results in their specific expansion in the *T. squamosa* genome. Interestingly, majority of *Tridacna* specific expanded POTs are tissue-specifically expressed in the symbiotic mantle (**Figure 3B**), which provides an alternative and efficient way of nitrogen acquisition through uptake of oligopeptides. Thus, it seems plausible that activation of symbiosis-dependent transcriptional programs operates in conjunction with a few sets of gene expansions to contribute to nitrogen-related metabolites uptake in the host giant clam.

In the photosymbioic holobiont, nitrogen-cycling is of fundamental importance to stability and resilience of the host, in particular under circumstances of nutrient deficiency in marine waters^44^. Previously, it has been proposed that external nitrogen (for example, in the form of NO_3_^-^) could be absorbed from the external seawater and fixed in ctenidial (gills), and then exported through the SCIAIN system to facilitate photosynthesis in symbiotic algae ^45^, an idea currently supported by our omics data. Moreover, our results also revealed that various protein hydrolytic enzymes involved in urea cycling pathways including CPD (Carboxypeptidase D), proteases, and peptideases, along with aminomethyltransferase (AMT) may co-participate in nitrogen cycling of giant clam-dinoflagellates symbiosis (**Figure 3C**). Therefore, dual NO_3_ ^-^ or ammonia host exporting pathways may guarantee a minimalist nitrogen supply to photosynthesizing symbionts, though the dynamic alterations in metabolite flux in response to climate change call for future investigation.

### Section 4: Light-sensing

As sunlight is the principal source of energy in many photosymbiotic organisms, the host has to evolve a light-sensing ability to ensure maximal or sufficient photosynthesis efficiency of symbiotic dinoflagellates. Anatomic evidence confirmed that a row of photoreceptors is located along the margin of the siphonal mantle (**Figure 4A**). Previously, cellular electrophysiological analysis have proved its functional relevance on the basis of their light-dependent hyperpolarization in the Tridacna giant clam ^46, 47^. Moreover, intracellular electrophysiological recordings have shown that the photoreceptors in the giant clam have distinct sensitivity peaks in response to specific wavelengths (that is, an ability of color discrimination), including green, blue and even ultraviolet regions ^47^. However, the mechanisms therein remain unelucidated.

**Figure 4.**
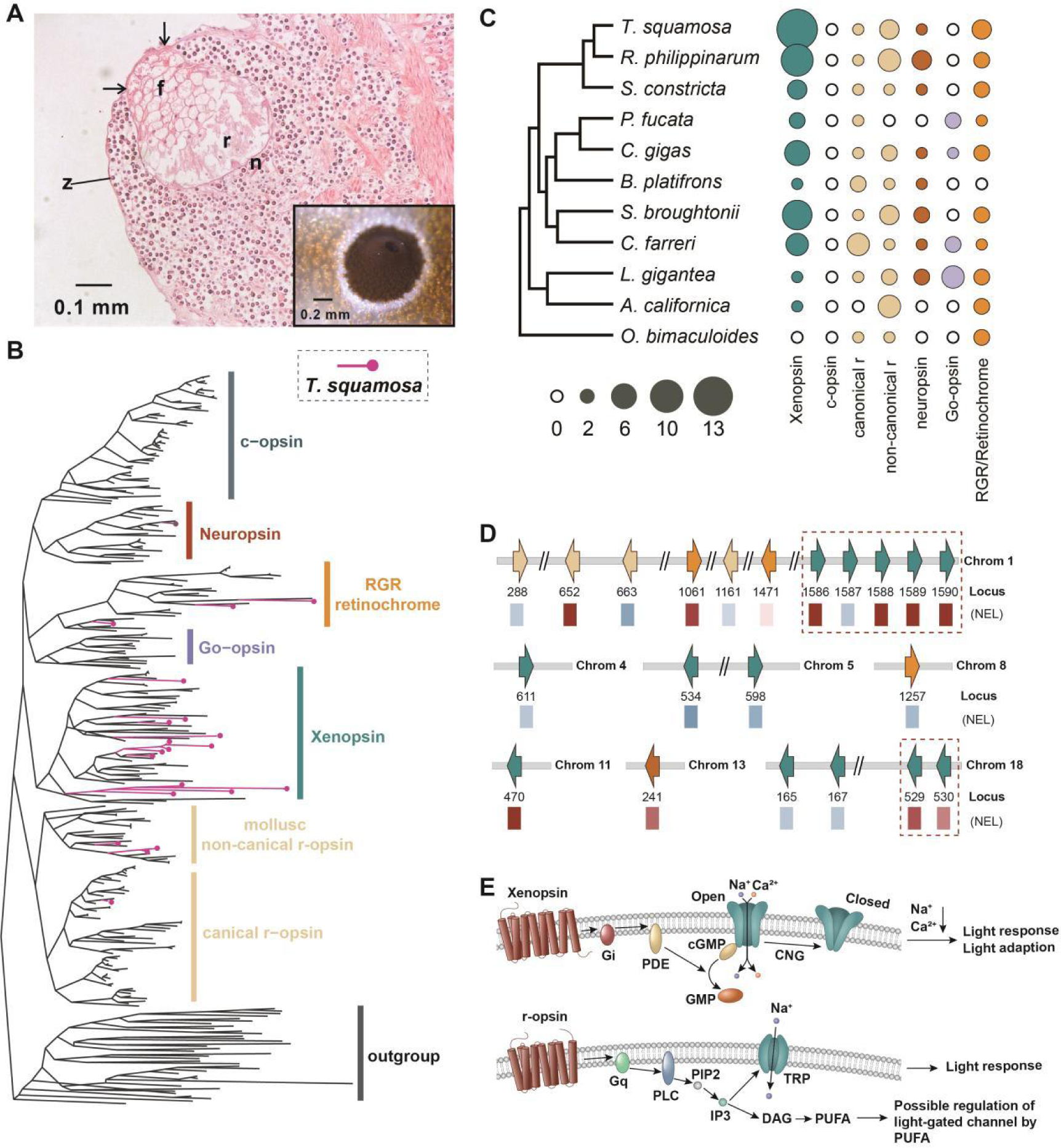
Photosensitive system in *T. squamosa* for light-sensing. **A,** Histological image of *T. squamosa* eyespots. Eyespots of clam is surrounded by *Symbiodinium spp.* zooxanthellae (z). Section through a single eye showing the positions of the Photoreceptor cells (r), “filler” cells (f), nerve bundle (n), and the limits of the pigmented aperture (arrowheads). **B,** Phylogenetic analysis of opsin superfamily from representaive metazoans, which is constructed using FastTree and visualized by FigTree. Sequences from *T. squamosa* genome are displayed in red color. **C,** Summary of known opsin complements within the molluscs. Bubble size indicates the genes number of the opsin paralogs, while its color distinguishes different opsin paralogs. **D,** A model for chromosomal organization and relative expression level of opsins genes. Opsin paralogs are indicated in colors as the same as C diagram. NEL, normalized expression level. **E,** Opsin signalling cascades in *T. squamosa*. PKC, protein kinase C; PLC, phosphoinositide-specific phospholipase C; PDE, phosphodiesterase; TRP, transient receptor potential channel; CNG, Cyclic nucleotide-gated channel.

Phylogenetic evidence showed that the *T. squamosa* genome encodes a total of 21 opsin genes with various diversified opsin types, including 13 xenopsin and 4 common r-opsin, 1 neuropsin and 3 RGR retinochrome, representing the maximum opsin gene number among known molluscans (**Figure 4B & 4C, supplementary Table 16**). In vertebrates, color discrimination is generally mediated by means of ciliary opsins (c-opsin) and its associated cascades ^48, 49^. However, c-opsin is absent in the *T. squamosa* genome, whereas 13 copies of xenopsins are present, which constitutes a lineage-specific expansion and highest number of genes in the single molluscan species (**Figure 4C, supplementary Table 16**). Interestingly, our comparison of the *Tridacna* xenopsins with the ciliary opsins of other species shows a certain degree of similarity in sequence and conserved motif structure, which distinguishes them from another common rhabdomeric opsin (r-opsin) (**Figure S21**). Therefore, it seems reasonable to assume that xenopsin, instead of c-opsin, may be functionally important in color sensitivity in protostomes lineages including the giant clam ^50^. In particular, five paralogs of the xenopsin subfamily are tandemly localized on a short region of chromosome 1 (71288623 to 71378942), and 4 of which exhibit a siphonal mantle-specific expression (**Figure 4D & supplementary Figure S22**), forming a light-sensing locus in *T. squamosa* genome. Moreover, expression patterns analysis revealed that both xenopsin and r-opsin and their associated signaling cascades are predominantly expressed in the symbiotic tissue, siphonal mantle (**Figure 4E, Figure S23**). Therefore, development of a light sensing locus via lineage-specific opsin expansion and co-existence of two active phototransduction pathway are likely crucial to the evolution of sophisticated light sensing and adaption to photosymbiotic lifestyle in the giant clam.

### Section 5: Photo-protection

Giant clams living in seawater require exposure to intense solar light that subserves photosymbiosis. As a price, host is also susceptible to detrimental photodamage due to inductive DNA mutation under ultraviolet radiation (UVR) in the solar light spectrum. In order to avoid such damage, the host has to develop a protective or repair system against solar radiation, which mechanisms have yet lacked substantiation. In our study, genomic and evolutionary evidence revealed that 6 of the photolyase/cryptochrome family (CPF family) genes are encoded in the *T. squamosa* genome, which are clustered into five subfamilies, namely: animal CRY, (6-4) PHR, CPD class II photolyase (PHR), cyclobutane pyrimidine dimers (CPDs) class III PHR, and CRY-DASH (**Figure 5A, Figure S24**). Among these, the (6-4) PHR, CPD class II PHR, and plant PHR subfamilies function as photolyases that respond to ultraviolet-induced DNA damage, while the others are chiefly responsible for light sensing in circadian rhythms regulation. It is worth noting that the 3 photolyase genes of *T. squamosa*, TsEVM0014143, TsEVM0019824, and TsEVM0003826, share a tissue-specific expression pattern in the siphonal mantle (**Figure 5A**). Additionally, all of the three photolyase genes retain conserved domains and sequence motifs of DNA photolyase and flavin adenine dinucleotide (FAD) binding, suggestive of repair function in damaged DNA binding and light energy utilization (**Figure S25**). In contrast, photolyase orthologs of non-symbiotic bivalve Crassostrea oysters do not exhibit mantle-specific expression patterns (**Figure S26**), further underscoring the significance of lineage-specific innovations for transcriptional regulation of a light-dependent DNA repair machinery that facilitates photosymbiotic adaptation in the *Tridacna* giant clam.

**Figure 5.**
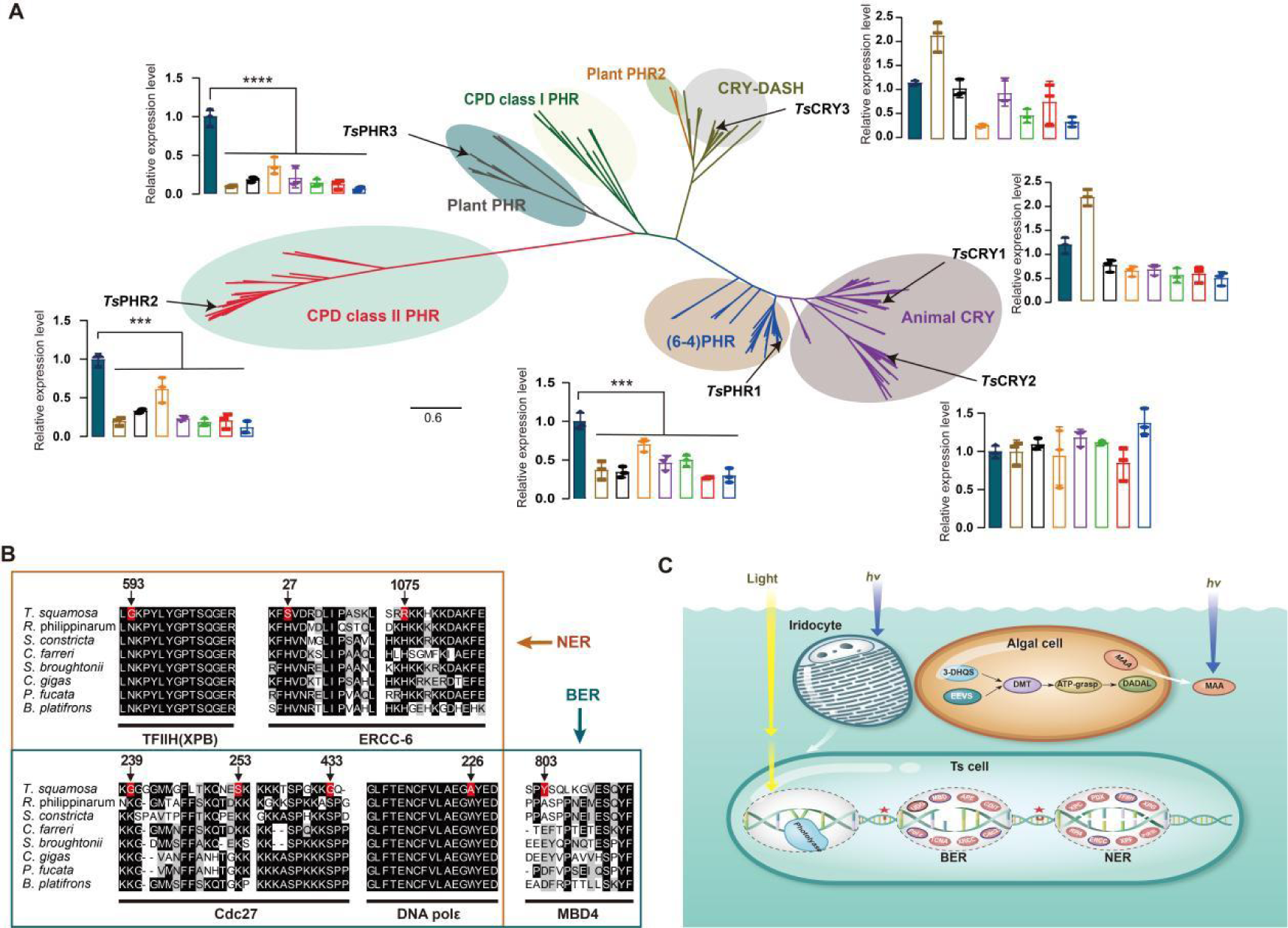
Photolyase and photosymbiosis-DNA repair governs photo-protection. **A,** Phylogenetic tree of photolyase/cryptochrome family (CPF family) in T. squamosa and other representative species. The CPF subfamilies are group in different color, and expression data of CPF genes in *T. squamosa* are shown beside arranged by siphonal mantle, adductor muscle, inner mantle, hemocytes, heart, ctenidium, foot, and hepatopancreas as as mean ± SE. *p <0.05, **p <0.01, and ***p<0.001. **B,** Positive selection analysis of core members of NER and BER. Eight closest species include T. squamosa was assigned to examine the selective constraints. Amino acids marked in red are positive selection sites for *T. squamosa*, *p < 0.05. **C,** Schematic summary of mechanisms for photo-protection in *T. squamosa*, including MAA produced by symbiotic algae, photosymbiosis-DNA repair, and iridocyte. Black circle marks the expanded gene family (UDG), and blue circle for positively selected genes (PSGs).

Furthermore, light-independent repair mechanisms (LIR) are widely present in many organisms, which includes nucleotide excision repair (NER) and base excision repair (BER) via involvement in the action of different glycosylases. Among them, uracil-DNA glycosylase (UDG) is one of the evolutionarily conserved glycosylases that can initiate base excision repair by removing uracil from DNA. Seven of the UDGs was found to be encoded and has apparently undergone lineage-specific expansion in the Tridacna genome (**Figure S27**), suggesting that both light-dependent and light-independent repair systems cooperate in the DNA damage repair in Tridacna species. Strikingly, positive selection was identified as a critical driving force for promoting diversity of functions and adaptive divergence^51, 52^. This idea was well supported by our evidence that multi-selective sites were present in the core members of NER and BER machinery, including DNA excision repair protein ERCC-6, TFIIH basal transcription factor complex helicase XPB subunit, DNA polymerase subunit Cdc27, DNA polymerase alpha/epsilon subunit B, and methyl-CpG-binding domain protein 4 (**Figure 5B**). Therefore, strong selective pressure likely provides a decisive driving force toward rapid adaptation by the DNA repair machinery in manners conducive to photosymbiotic lifestyle in the giant clams.

In addition, *Tridacna* giant clams have also evolved multiple complementary strategies to defend against harmful UVR. To illustrate, UV photoreceptors in the giant clam exhibits UV sensitivity, and can trigger reflex behavior of defensive withdrawal when under overwhelming radiation exposure ^47^, and closure of the opaque shell for physical protection against UV radiation. The giant clam has also evolved an alternative photo-protection mechanism by the generation of iridocytes within mantle, which can prevent photodamage through absorption of potentially harmful UVR and scatter light at longer wavelengths simultaneously (**Figure 5C**). However, how iridocyte differentiation is achieved requires mechanistic clarification in the future. In addition to the host, symbiotic dinoflagellates also contribute directly to photo-protective defenses in the host. Specifically, mycosporine-like amino acids (MAAs) are potent UV-absorbing compounds, which are supposed to be produced in the algal symbionts and deposited in the host mantle ^53^. Our genomic evidence corroborated this assumption. We confirmed this by observation of a lack of MAAs biosynthetic pathways in the *T. squamosa* genome, but the presence of key biosynthetic genes in the transcriptome of the symbiont, which further illustrates the mutually beneficial nature of photosymbiosis in the giant clam-dinoflagellates pair (**Figure 5C**).

## Conclusion

As a prominent member of marine habitats in coral reefs, the fluted giant clam deserves special attention, ecologically and evolutionarily. Not only does it provide shelter and nutrient sources to cohabitant species of the reef environment, but also serves as an important, albeit underappreciated, booster of biodiversity by being an integral part of carbon-nitrogen cycles in the coral reef landscape. Indeed, much of the known ramifications of climate change such as global warming and ocean acidification also afflict *T. squamosa*, with striking resemblances to maladies seen in corals. This collaterality can in part be accounted for by the fact that both corals and giant clams reply heavily on exquisitely regulated symbiotic systems for maintaining integrity and health. In the case of *T. squamosa*, failure for juvenile larvae to acquire dinoflagellates in a critical developmental window leads to metabolic incompetence and lethality. Even for adults, inopportune conditions such as warming seawater can cause the expulsion of zooxanthellae and subsequently catastrophic bleaching, thus severing the photosymbiotic bond. In this study, we have provided a comprehensive genomic framework for delineating and describing prototypic photosymbiosis in the giant clam-dinoflagellates pair. In particular, we highlighted rapid and expansive transcriptional changes and tissue-specific gene expressionsin the host that are apparently instrumental to metabolic remodeling following productive establishment of symbiosis. Adding fresh nuances to photosymbiotic regulation is the multi-omic evidence that dinoflagellates actively contribute to host health by facilitating macromolecular repair against solar irradiation. Arguably, these observations may, by extrapolation, provide some modest inspirations to studies on reciprocally beneficial symbiotic interactions between microbiota and higher organisms. Collectively, our work here has advanced a fuller picture on the evolution and molecular features of marine photosymbiosis, which hopefully helps inform more focused biological investigation on preserving biodiversity and saving the environment.

## Methods

Three fluted giant clams (Tridacna squamosa) were collected from the Xisha Islands in the South China Sea during a biological resources reconnaissance survey. Adductor muscle tissue was dissected from the clams to extract DNA for Illumina or PacBio sequencing. Illumina sequencing involved the construction of paired-end libraries and subsequent sequencing on an Illumina HiSeq 2500 platform. The obtained raw reads were processed to remove sequencing adaptors and filter out contaminated and low-quality reads ^1^. For PacBio sequencing, genomic DNA was sheared and processed for single-molecule real-time (SMRT) bell preparation, followed by sequencing on a PacBio RS II platform with C4 chemistry. The resulting subreads were used for genome assembly.

The error-corrected subreads from both sequencing methods were utilized for draft assembly using specific software modules such as Canu v.1.5 ^2^, LoRDEC v.0.6 ^3^, and wtdbg (https://github.com/ruanjue/wtdbg). Genome quality was assessed by mapping Illumina reads, verifying genome completeness using conserved eukaryotic genes and metazoan benchmarking universal single-copy orthologues by using BUSCO v.3.0.2b ^4^, and performing Hi-C sequencing and assembly by HiC-Prov2.8.1 ^5^ and LACHESIS ^6, 7^. The genome was annotated for transposon elements (TE) and tandem repeats using a combination of homology-based and de novo approaches by RECON (version 1.08) ^8^ and RepeatScout ^9^. Protein-coding genes were predicted based on various methods including homologue, RNA sequencing (RNA-seq), and de novo gene prediction techniques, including PASA v.2.0.2 ^10^, TransDecoder v.2.0 ^11^and GeneMarkS-T v.5.1 ^12^. Predicted genes were further annotated against annotated against the Pfam database of the HMMER v.3.1b2 software (http://www.hmmer.org) and the InterPro database of InterProScan v.5.34-73.0 (https://github.com/ebi-pf-team/interproscan).

The protein-coding sequences in the *T. squamosa* genome were compared with those of other species to cluster gene families and identify single-copy orthologous genes using OrthoFinder (v2.4.0) ^13^software. Phylogenetic relationships were analyzed with RAxML v.8.2.12 ^14^, and gene family expansion and contraction were determined based on species divergence time estimated using MCMCTree from the PAML package ^15^. Positive selection analysis was conducted to examine the selective constraints on candidate genes ^15^. Oxford dot plots were used to assess the conservation of macrosynteny between species with chromosome-level assemblies ^16^. Additionally, karyotype analysis was performed using trochophore larvae chromosome preparations and staining protocols ^17^.

RNA isolation, Illumina RNA-Seq library preparation, and transcriptomic analysis were carried out on various tissues and at different time points to evaluate gene expression dynamics and identify tissue-specific genes. The R/Bioconductor package MaSigPro ^18^ was applied to normalized data to identify gene expression dynamics across different time point (0, 12, 24, 36, 48, 72 hours). Weighted correlation network analysis (WGCNA) ^19^ was applied to analyze differential genes in different tissues and identify gene ontology and KEGG enrichments for each module. The eyespot from the siphonal mantle of the clam was processed for histological observation, involving fixation, embedding, sectioning, staining, and microscopic examination.

## Data availability

The giant clam genome project has been deposited with the NCBI under the BioProject number PRJNA673920. The whole-genome sequencing data were deposited with the sequence read archive (SRA) database under accession nos. SRR13022221–SRR13022222 for Illumina sequencing and SRR13022210, SRR13022177, SRR13022188, and SRR13022199 for PacBio sequencing. The RNA-seq data have also been deposited with the SRA database under accession nos. SRR13022154, SRR13022156-SRR13022165, SRR13022167-SRR13022176, and SRR13022178-SRR13022180 for tissue distribution, and SRR13022181-SRR13022187, SRR13022189-SRR13022198, and SRR13022200-SRR13022204 for time course symbiosis establishment. The Hi-C data has been deposited with the SRA database under accession nos. SRR13022166

