## supplementar figures for "Genomic insights into photosymbiotic evolution in *Tridacna squamosa*"

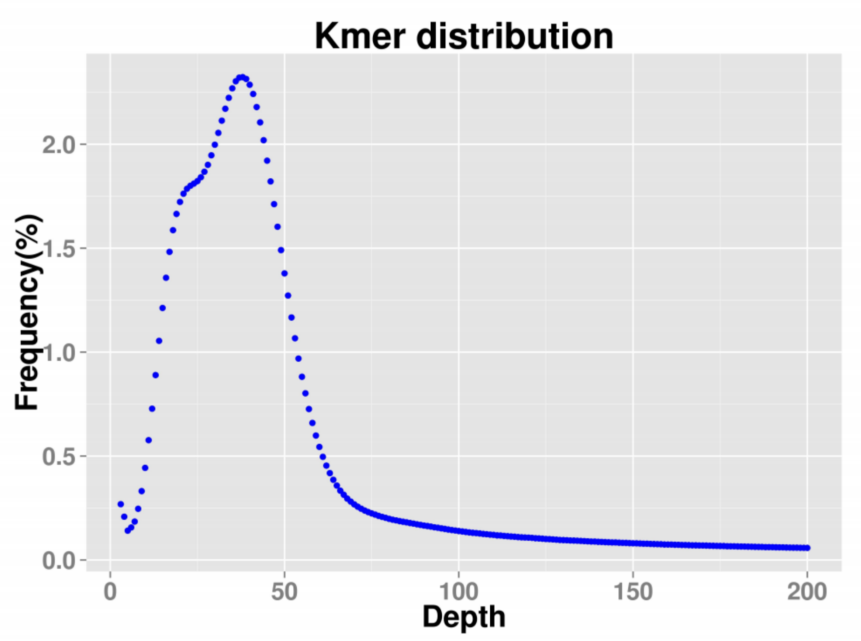


**Figure S1 k-mer distribution of the *Tridacna squamosa* genome.** Genome size estimation was performed by the k-mer analysis, and about 95.06 Gb corrected PacBio sequencing reads were selected to estimate the genome size. The genome size of T. squamosa thus estimated is 1.09 Gb.

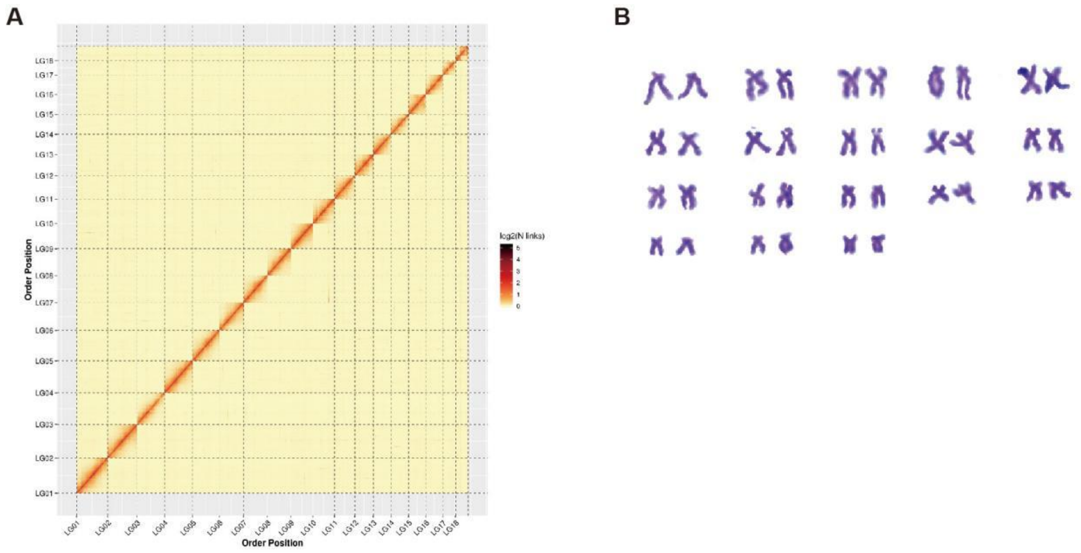


**Figure S2 Hi-C heatmap based on the chromosome-scale assembly of the *T. squamosa* genome.** A. The heatmap represents the contact matrices generated by aligning the Hi-C data to the chromosome-scale assembly of the *T. squamosa* genome. B. Conventional karyotype colored with Giemsa obtained from the *T. squamosa*.

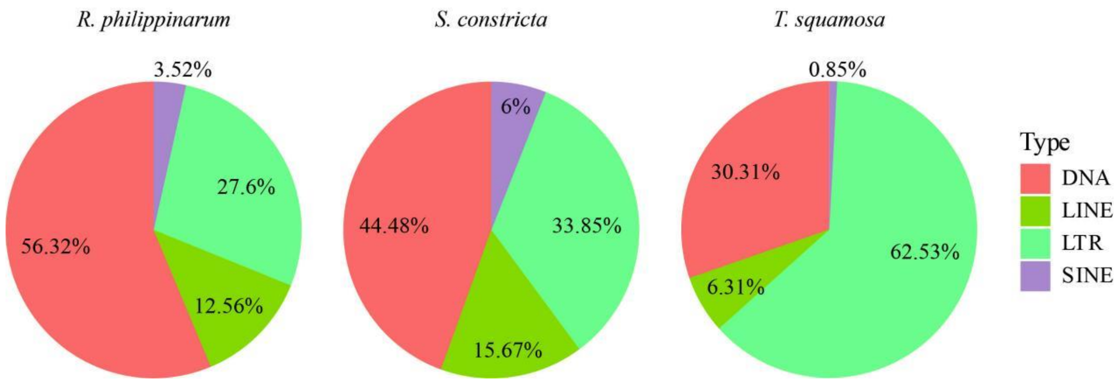


**Figure S3 Analyses of transposable elements in *T. squamosa* genomes.** Proportions of DNA transposons, LTR, LINE and SINE retrotransposons in the genomes of *T. squamosa* and its closed relatives *Ruditapes philippinarum* and *Sinonovacula constricta*

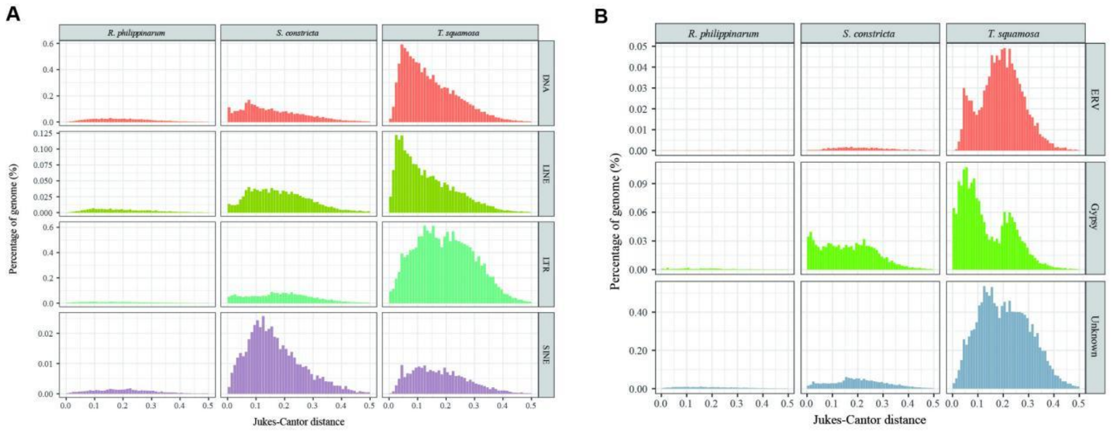


**Figure S4 Transposable element insertion history (Jukes-Cantor distance adjusted)**. The distribution of sequence divergence rates of TE (A) and LTR (B) as percentages of the genome size of *T. squamosa*, *R. philippinarum* and *S. constricta*

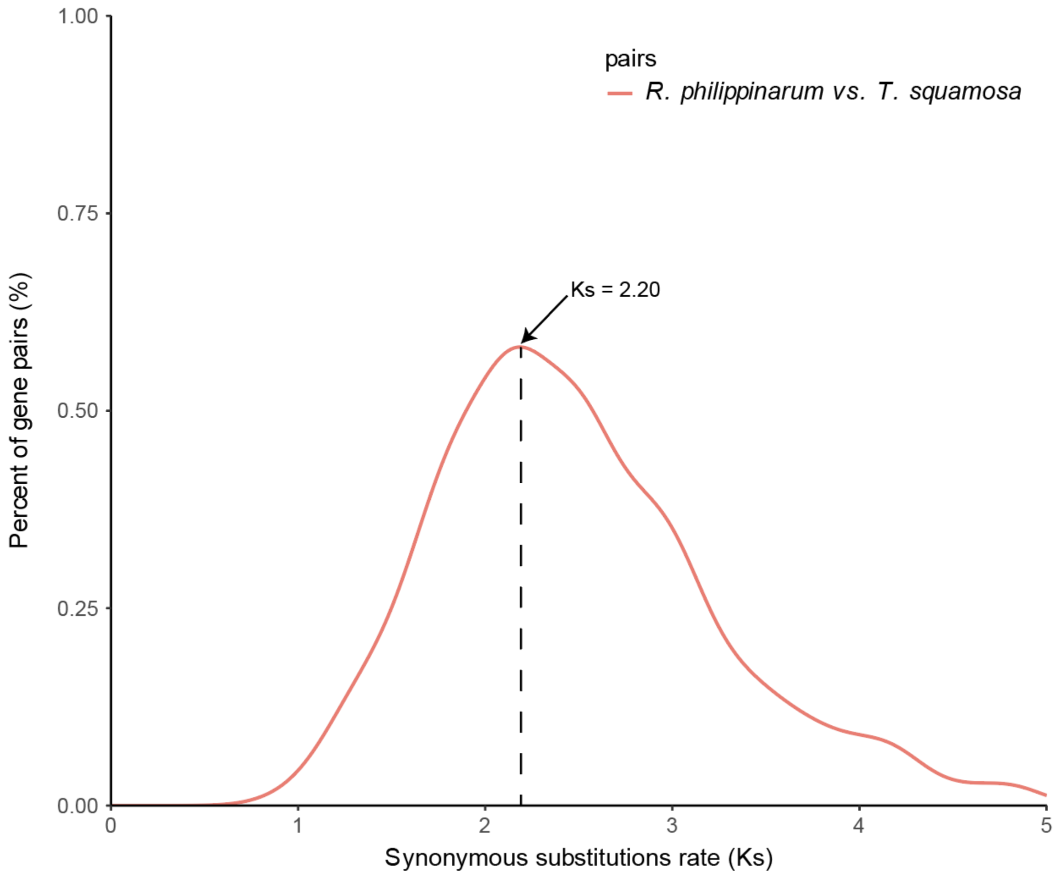


**Figure S5 Distribution of the estimated synonymous substitutions rate (Ks) between T. squamosa and the selected orthologous genes.** Peaks represent the divergence between *T. squamosa* and *R. philippinarum*.

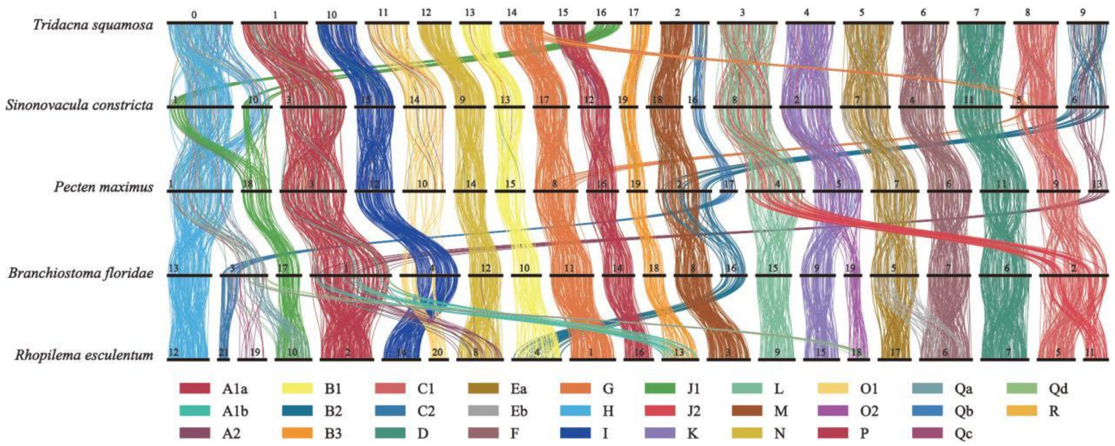


**Figure S6 Conserved syntenies.** Ribbon diagram showing conserved syntenies among *Tridacna squamosa (the giant clam)*, *Sinonovacula constricta (Chinese razor clam)*, *Pecten maximus (the giant scallop)*, the bilaterian amphioxus *Branchiostoma floridae*, and jellyfish *Rhopilema esculentum* ($\alpha \leq 0.05$, permutation test one-sided false-discovery rate). The vertical lines between species represent orthologous genes, coloured according to the BCnS synteny groups [1]. Only groups of genes that have significantly conserved chromosome-scale linkage (synteny) between metazoan species are shown.

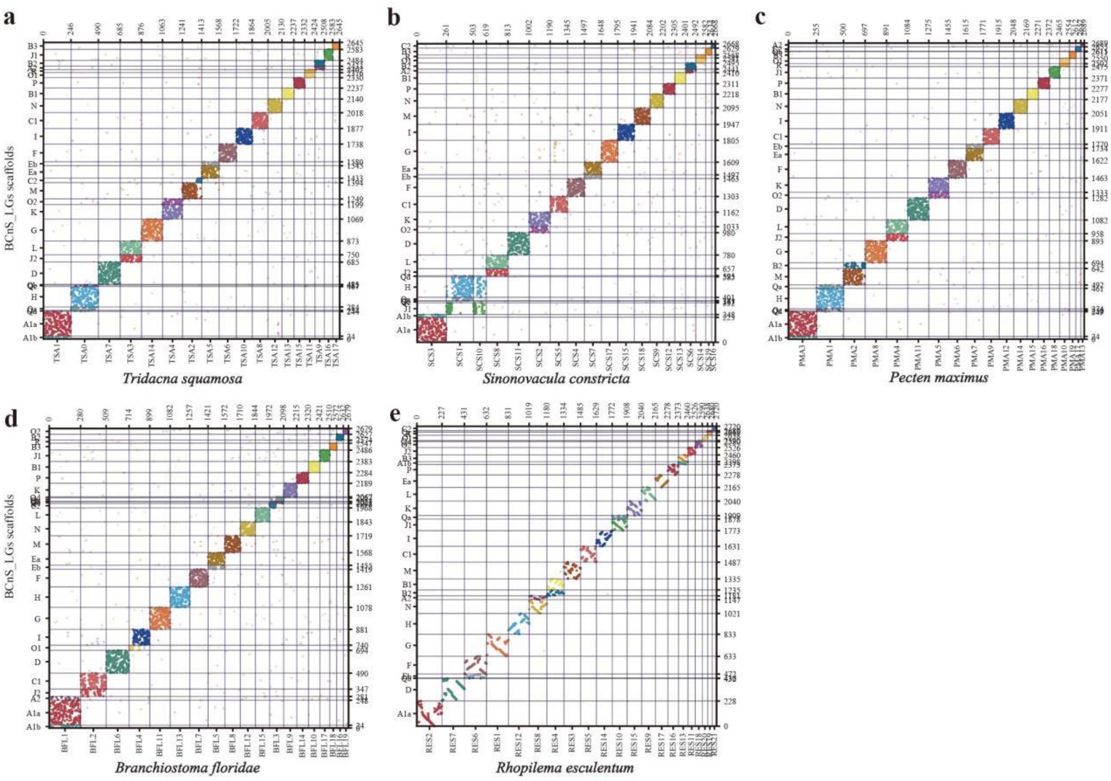


**Figure S7 Chromosome-based macrosynteny analysis reveals the high degree of conserved synteny in metazoa.** Chromosome-based macrosynteny is shown in the form of Oxford dot plots with comparisons between the chromosomes of *Tridacna squamosa* (this study), *Sinonovacula constricta*, *Pecten maximus*, *Branchiostoma floridae*, *Rhopilema esculentum* (*x* axis) and the previously reconstructed 29 BCnS-ALGs (*y* axis) [1].

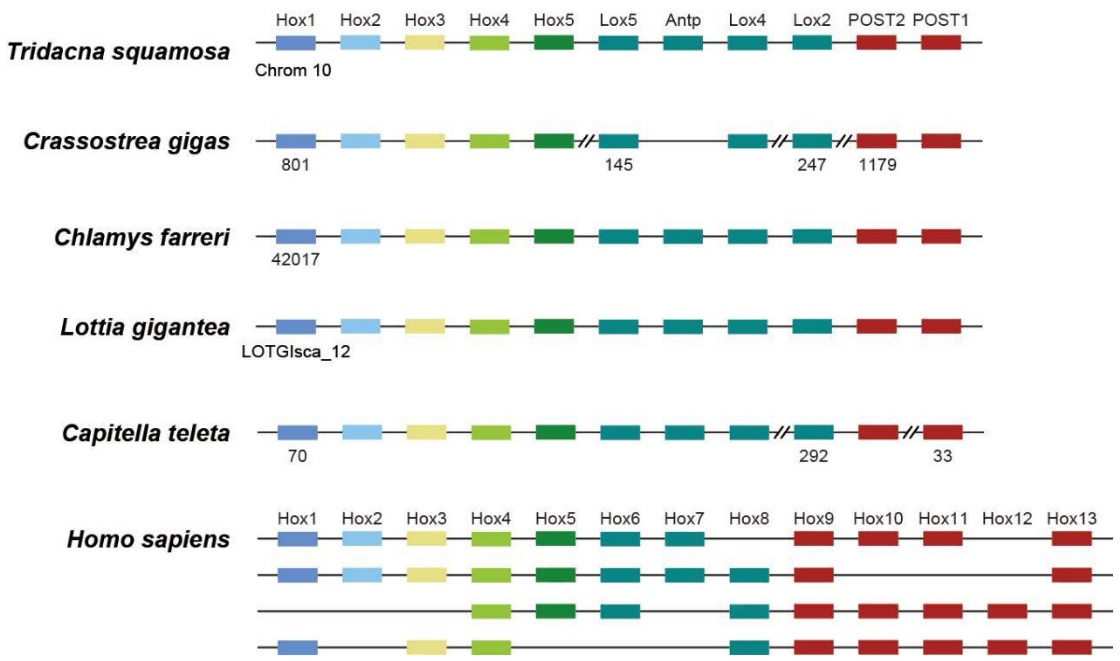


**Figure S8 Schematic representation of Hox gene clusters in metazoan genomes.** Comparison of chromosomal organization of Hox gene clusters of *Tridacna squamosa* with other animals. Different Hox genes are labelled with coloured boxes. Double slashes indicate that the scaffold of the Hox cluster is non-contiguous or interrupted.

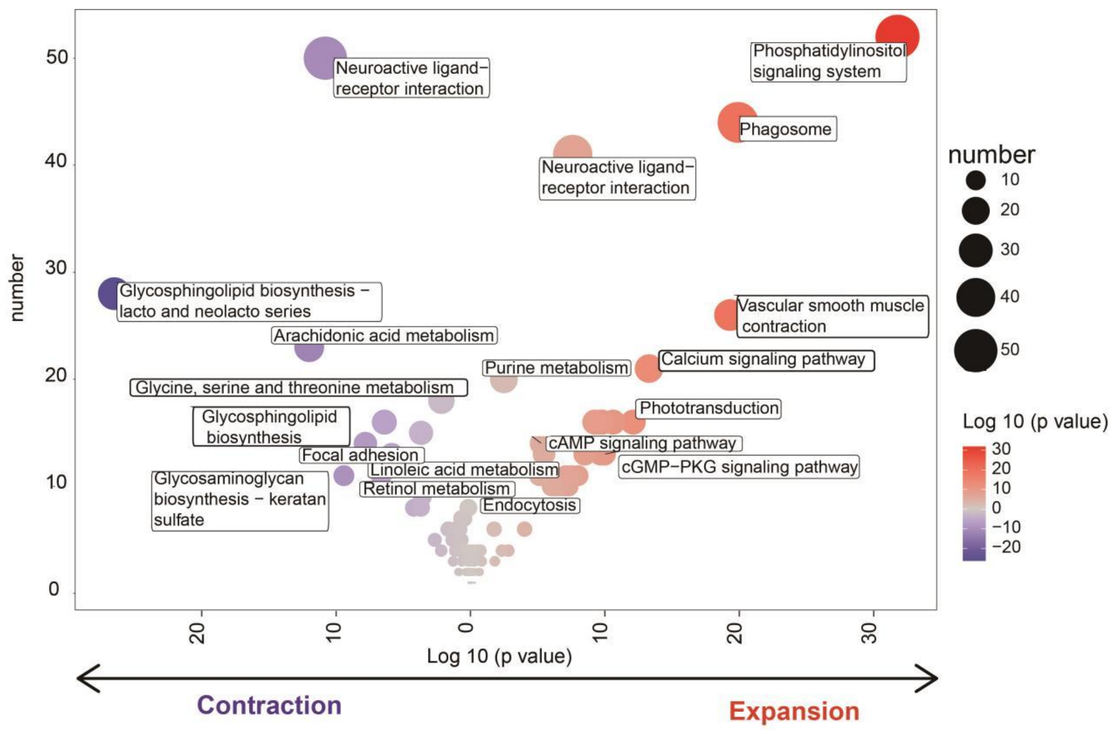


**Figure S9 KEGG enrichment analysis of the contracted and expanded gene family.** The size of the bubbles represents the number of genes enriched in the pathway, while the color of the bubbles represents the p-value. The purple color scheme indicates contracted gene families, while the red color scheme indicates expanded gene families.

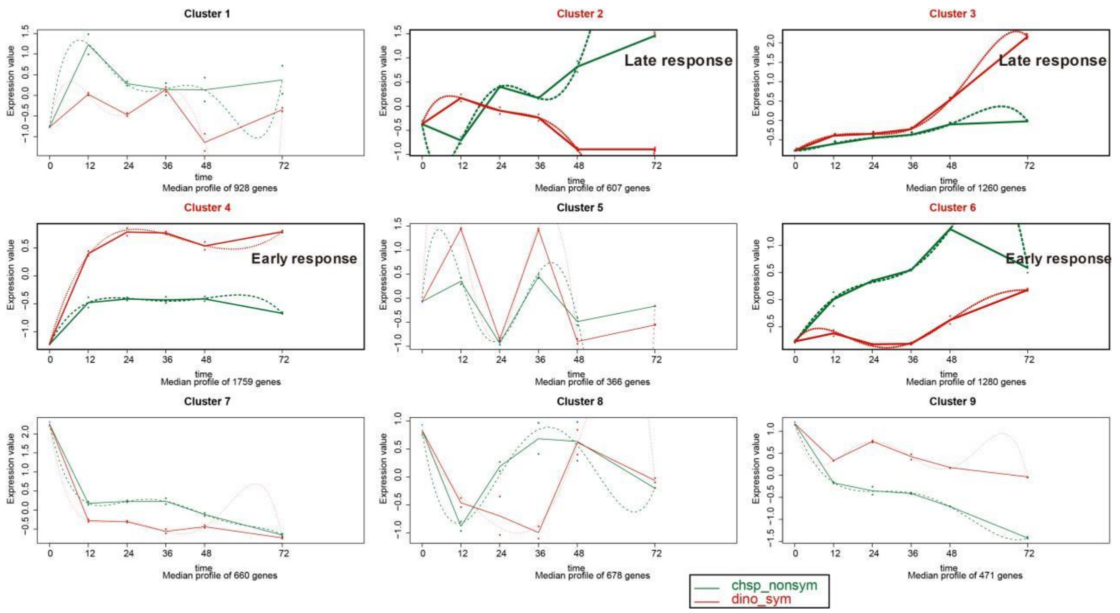


**Figure S10 MaSigPro clustering of RNA-Seq time series dataset from the symbiotic group and nonsymbiotic group.** The 9 clustered plots display time on the x-axis (algae feeding duration of 0-72 h) with relative expression on the y-axis. Each plot depicts the expression profile of the microarray probes within that cluster for dinoflagellates-feeding larvae (dino_sym, red) and chrysophyceae-feeding larvae (chsp_nonsym, green). Among them, four plots for further analysis are bolded, representing early and late symbiotic response, and early and late nonsymbiotic response. Different expressed genes were calculated as Q=0.05.

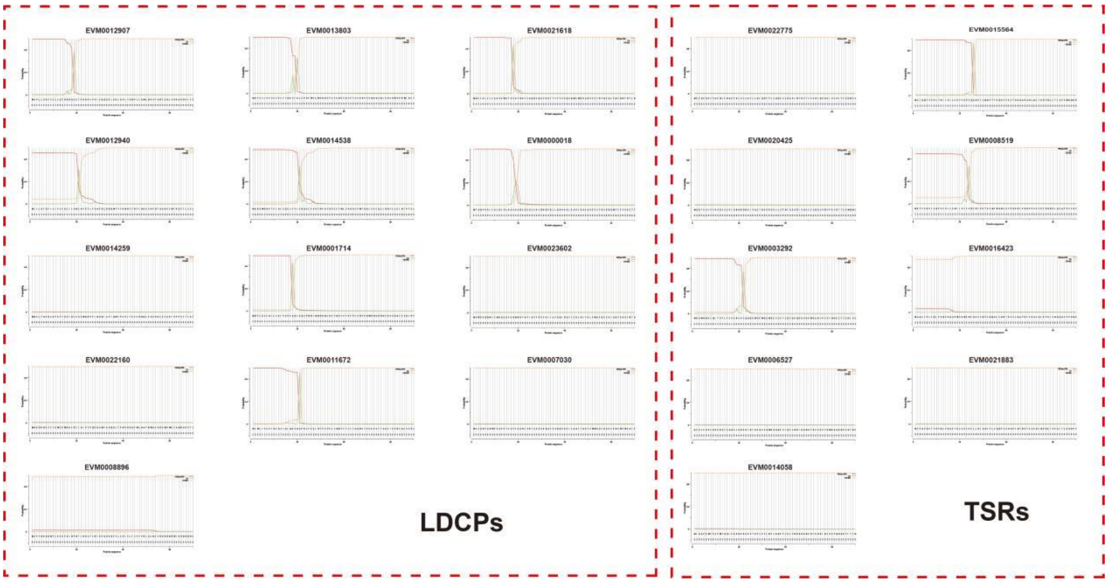


**Figure S11 Signal peptide analysis of pattern recognition receptors.** Signal peptide prediction of lectin domain containing proteins (LDCPs, left) and thrombospondin type 1 repeat (TSRs, right) using the SignalP 3.0 server according to SignalP Neural Networks (SignalP- NN).

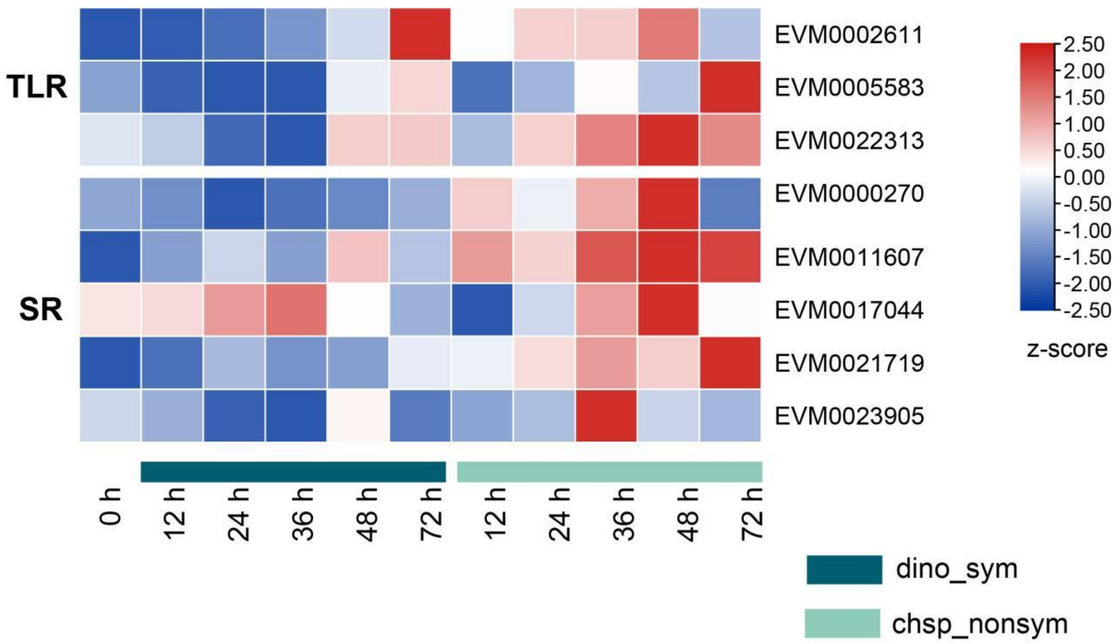


**Figure S12 Expression level of TLR and SR during algae feeding (0-72 h).** Heatmap shows the expression profile of Toll like receptors and scavenger receptors (SRs) at different time point in dinoflagellates-feeding larvae (dino_sym) and chrysophyceae-feeding larvae (chsp_nonsym). Colored bars represent z-score calculated from RPKM-values of a target gene in different tissues.

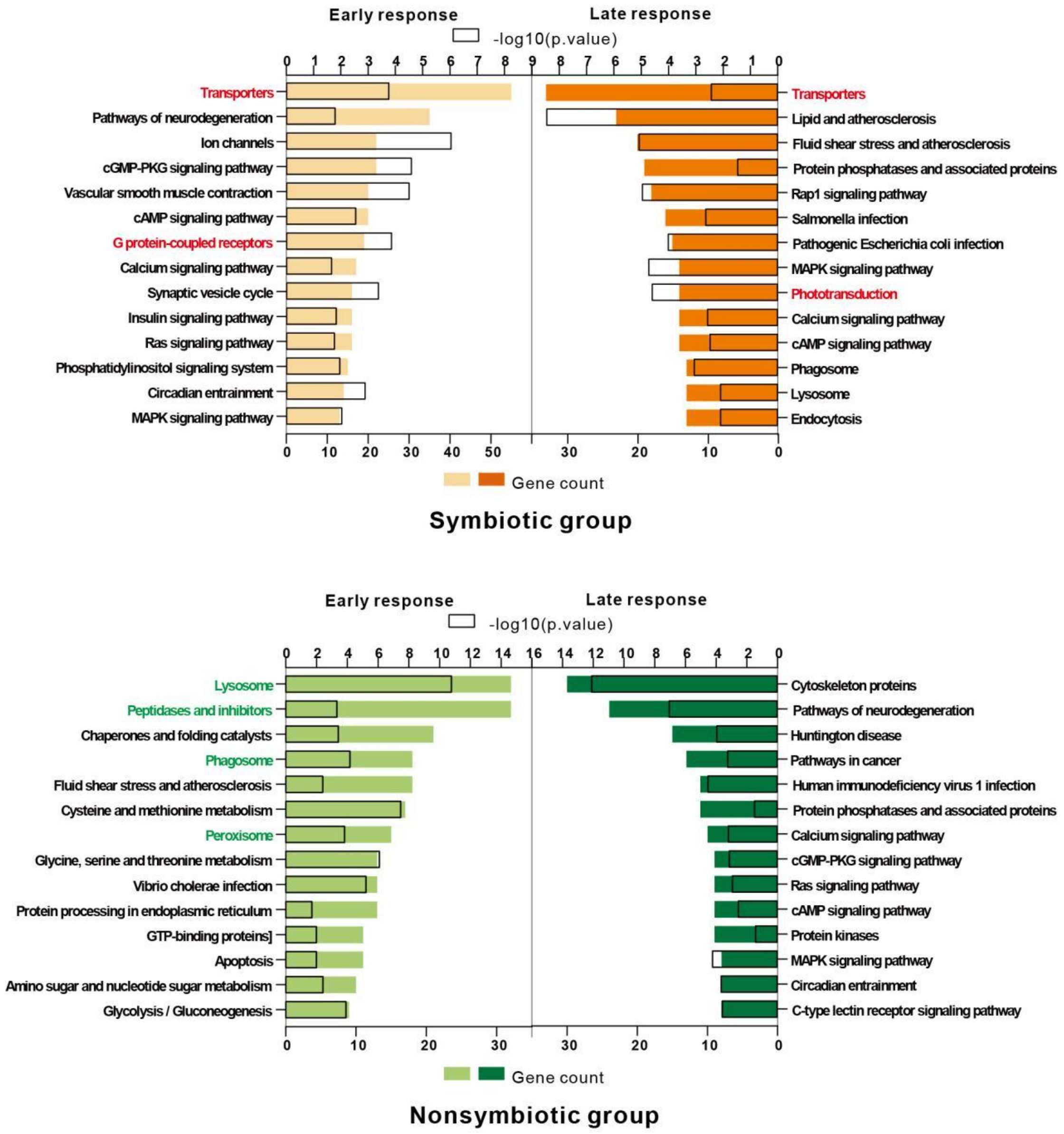


**Figure S13 KEGG enrichment analysis of gene sets in symbiotic group and nonsymbiotic group.** The gene count of different pathways is represented in color rectangle, while the p. value is shown in black box as -log10(p. value). Early response refers to the gene set that changed after 12 hours of algae-feeding, while late response refers to the gene set that changed after 48 or 72 hours

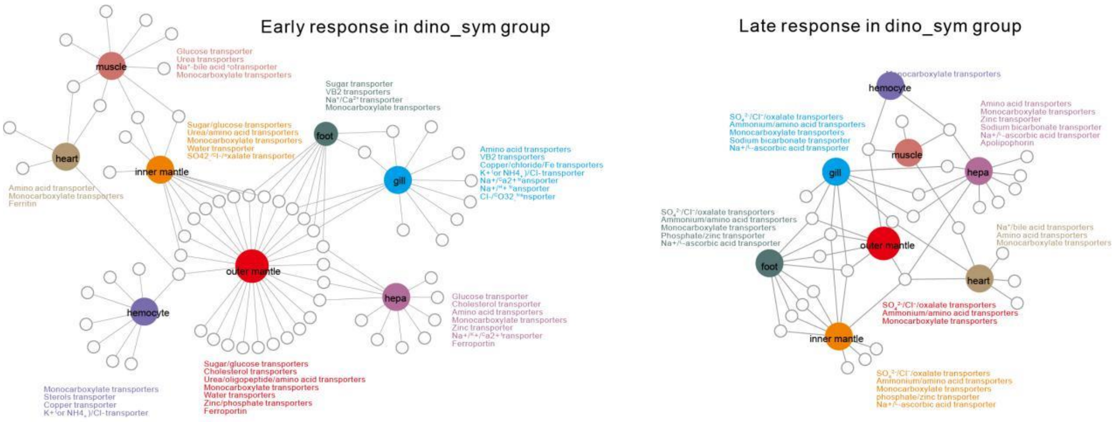


**Figure S14 Tissue distribution of deferentially expressed transports during symbiotic establishment.** Transporters identified in dinoflagellates-feeding (dino_sym) group were arranged for tissue correlation analysis using TissueEnrich packages (https://bioconductor.org/packages/release/bioc/html/TissueEnrich.html).

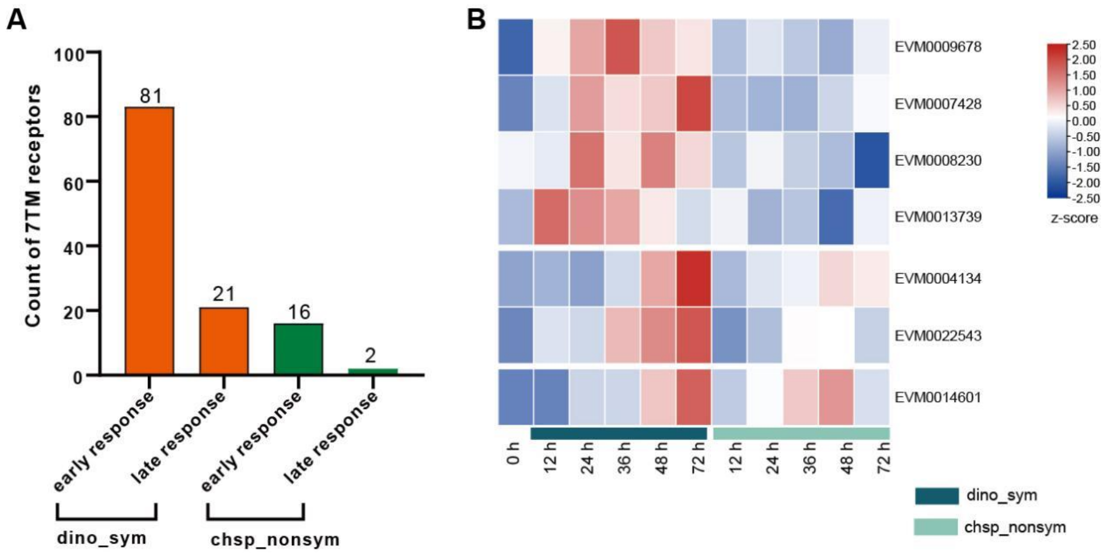


**Figure S15 Analysis of GPCR-arrestin cascade during algae-feeding (0-72 h)**. A. Number of 7TM receptors identified from DEGs in dino_sym group and chsp_nonsym group, respectively. B. Expression profile of arrestins at different time point in dinoflagellates-feeding larvae (dino_sym) and chrysophyceae-feeding larvae (chsp_nonsym). Colored bars represent z-score calculated from RPKM-values of a target gene in different tissues.

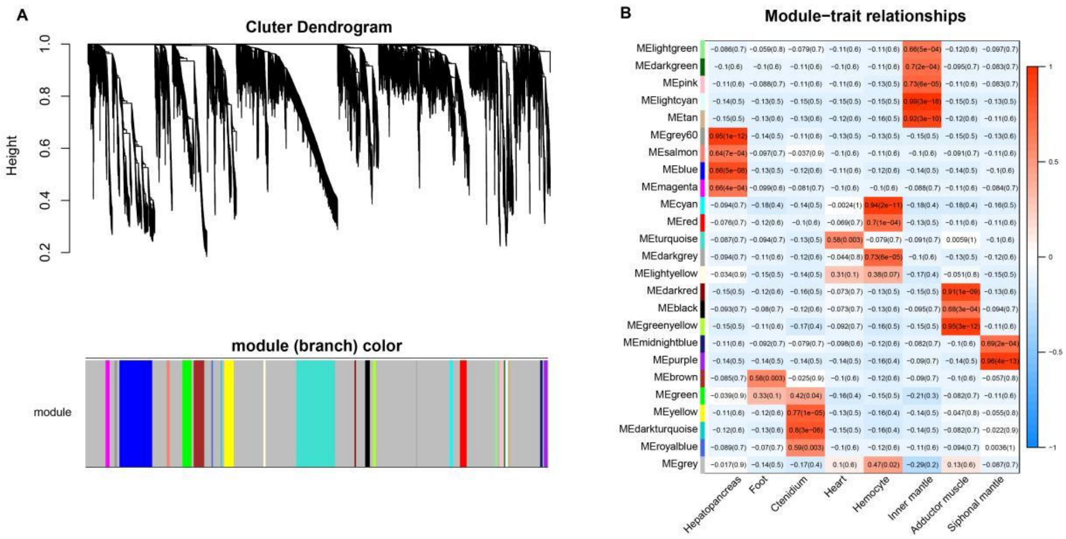


**Figure S16 The WGCNA analysis of RNA expression profile of different tissues.**

A. The cluster dendrogram of differentially expressed mRNAs. B. The module-trait relationship analysis between the 8 tissues. ME: module eigengenes.

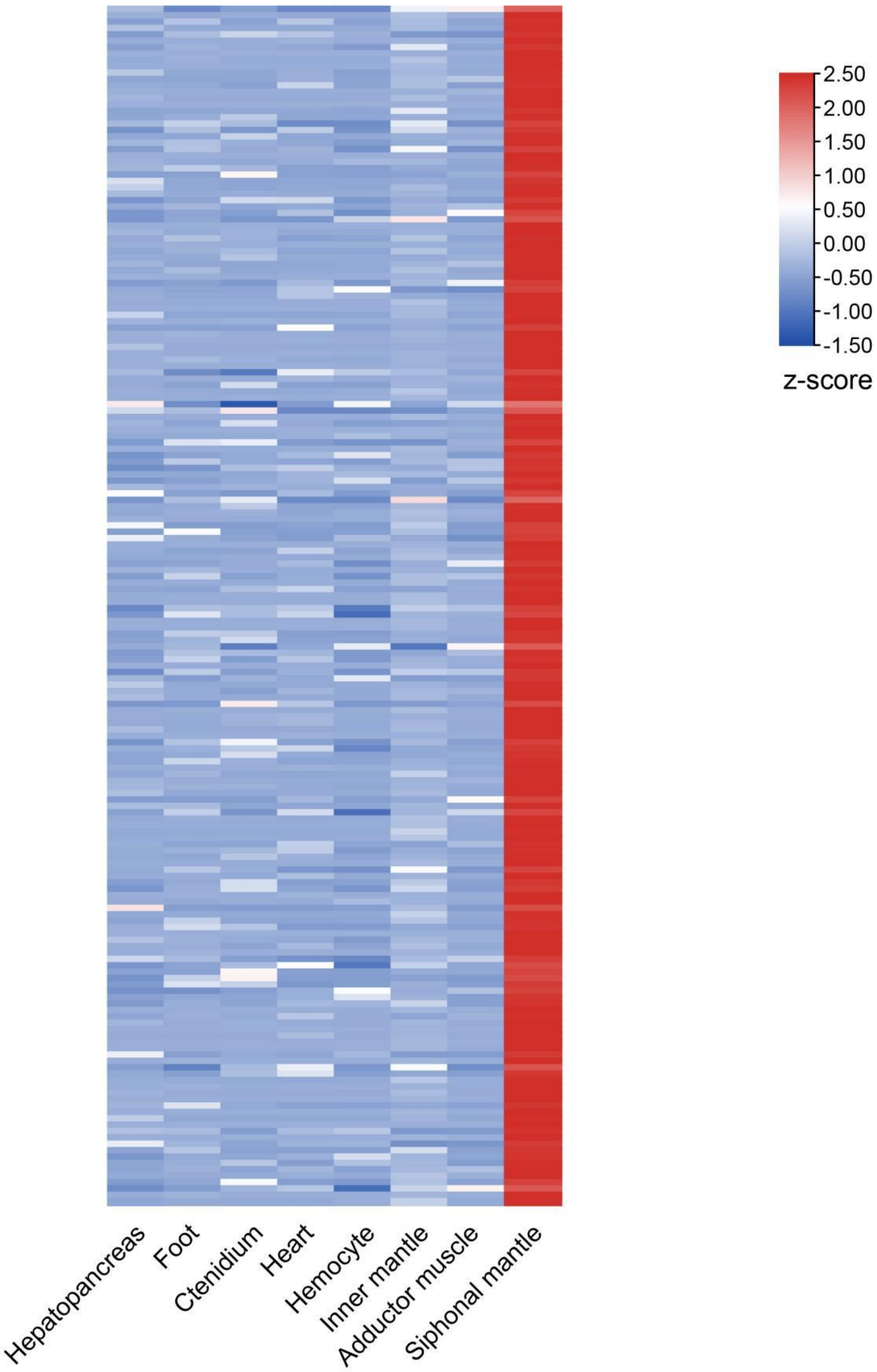


**Figure S17 Tissue distribution of 188 genes in modules significantly related to the outer mantle.** Heatmap shows the expression profile of 188 genes in genes in modules significantly related to the outer mantle, all of which specifically expressed in the outer mantle. Colored bars represent *z*-score calculated from RPKM-values of a target gene in different tissues.

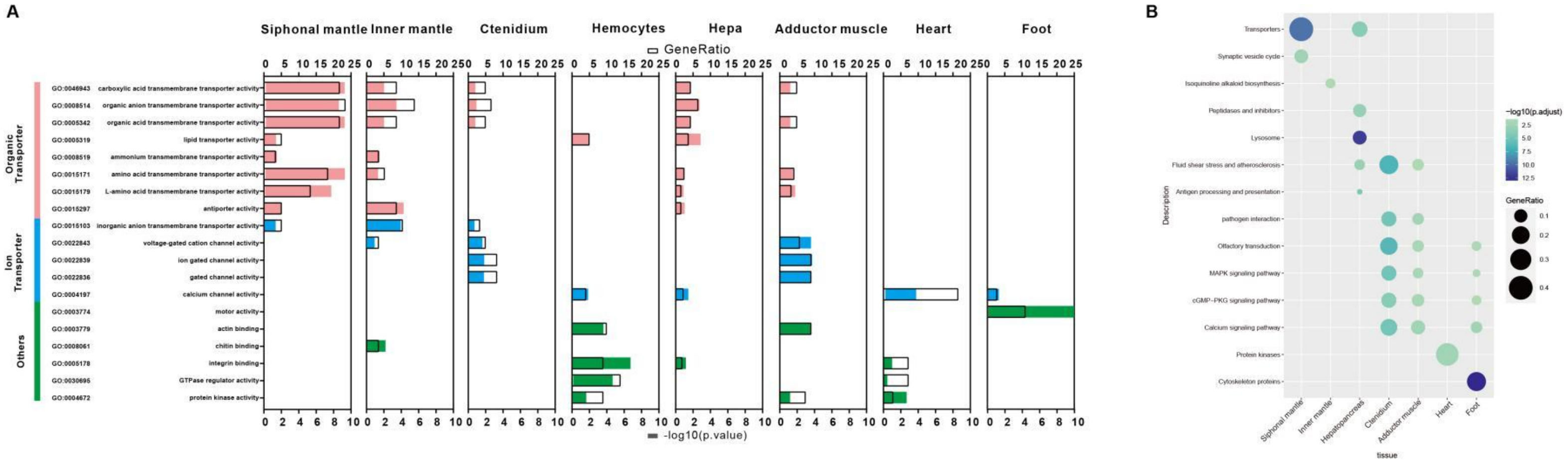


**Figure S18 Function enrichment analysis of RNA-seq datasets in different tissues by R package clusterProfiler.** A. Molecular function of Gene Ontology (GO) enrichment analysis.The Gene Ratio of different pathways is represented in color rectangle, while the p. value is shown in black box as -log10(p. value). B. KEGG pathway enrichment bubble plot. The size of the dots represents the gene ratio in the corresponding pathway, and the color indicates -log10(p. value)

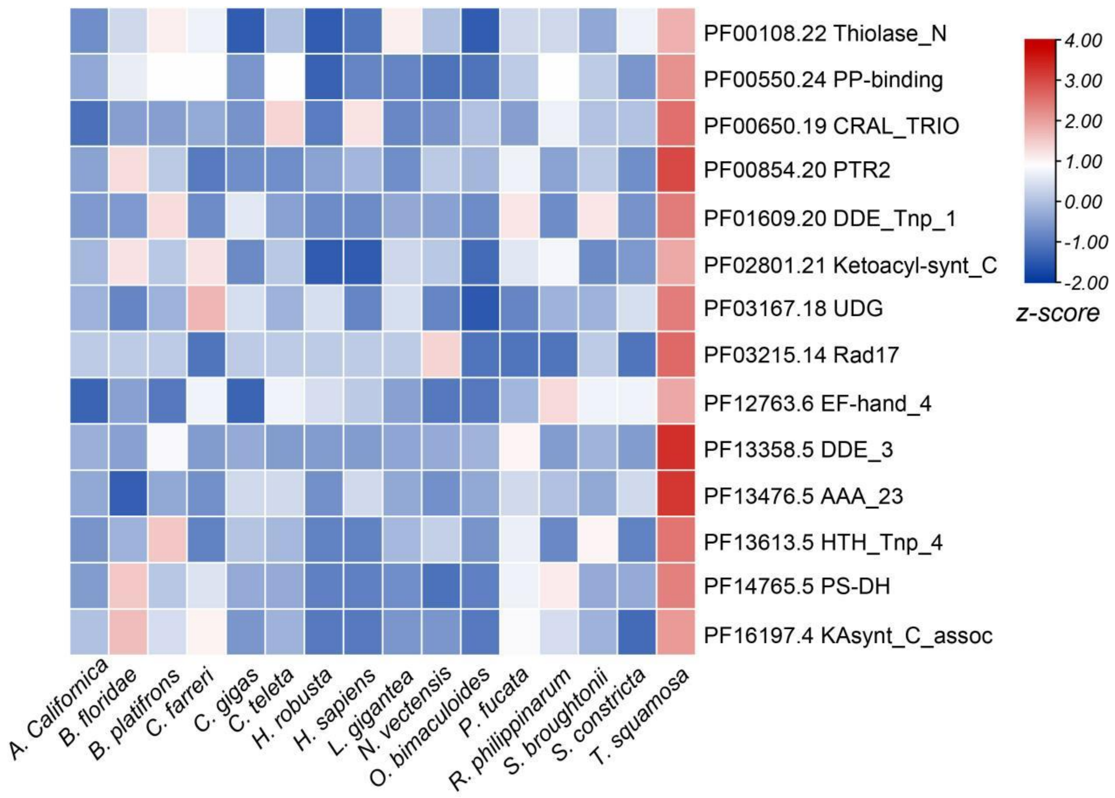


**Figure S19 Heatmap of expanded pfam family in *Tridacna squamosa* genome.**

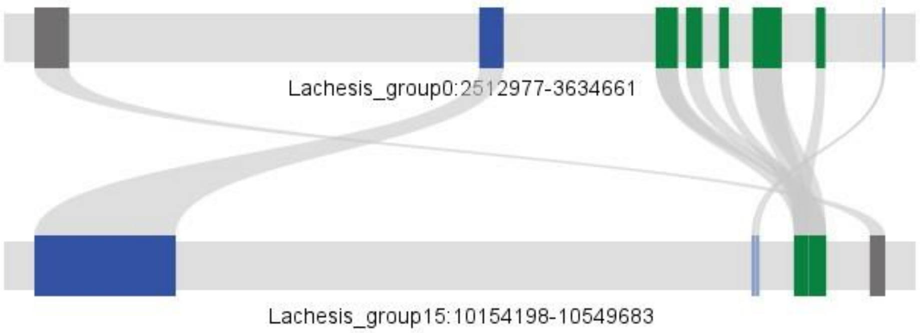


**Figure S20 Collinearity analysis of the POT gene family in *T. squamosa*.** The lines represent gene collinearity in the genome. Lachesis_group0, Chr1; Lachesis_group15, Chr14.

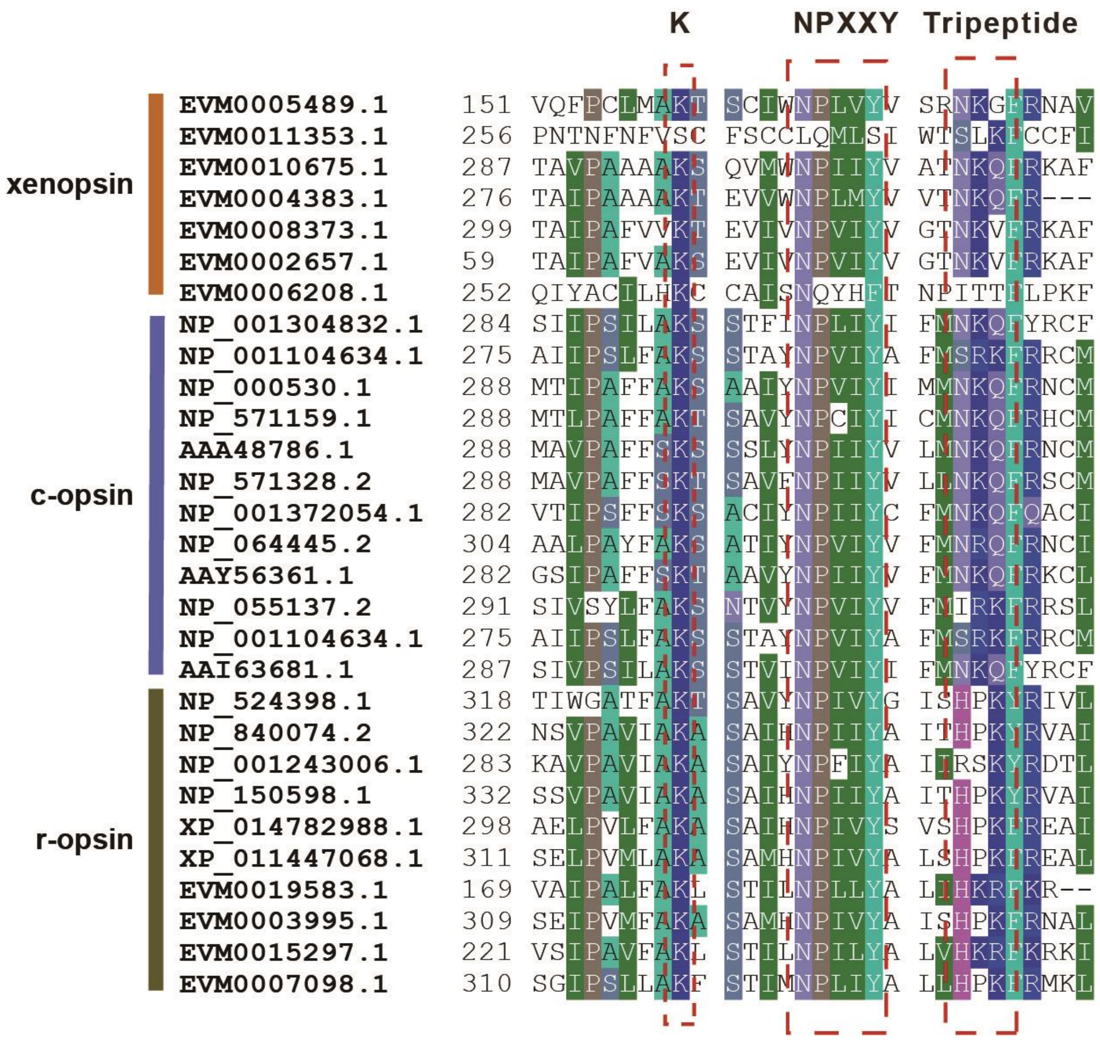


**Figure S21 Sequence alignment of *T. squamosa* xenopsin with c-opisn and r-opsin.** Sequence alignment was conducted and displayed using BioEdit Sequence Alignment Editor. The conserved motif (K, NPXXY, and tripeptide) are marked with red box.

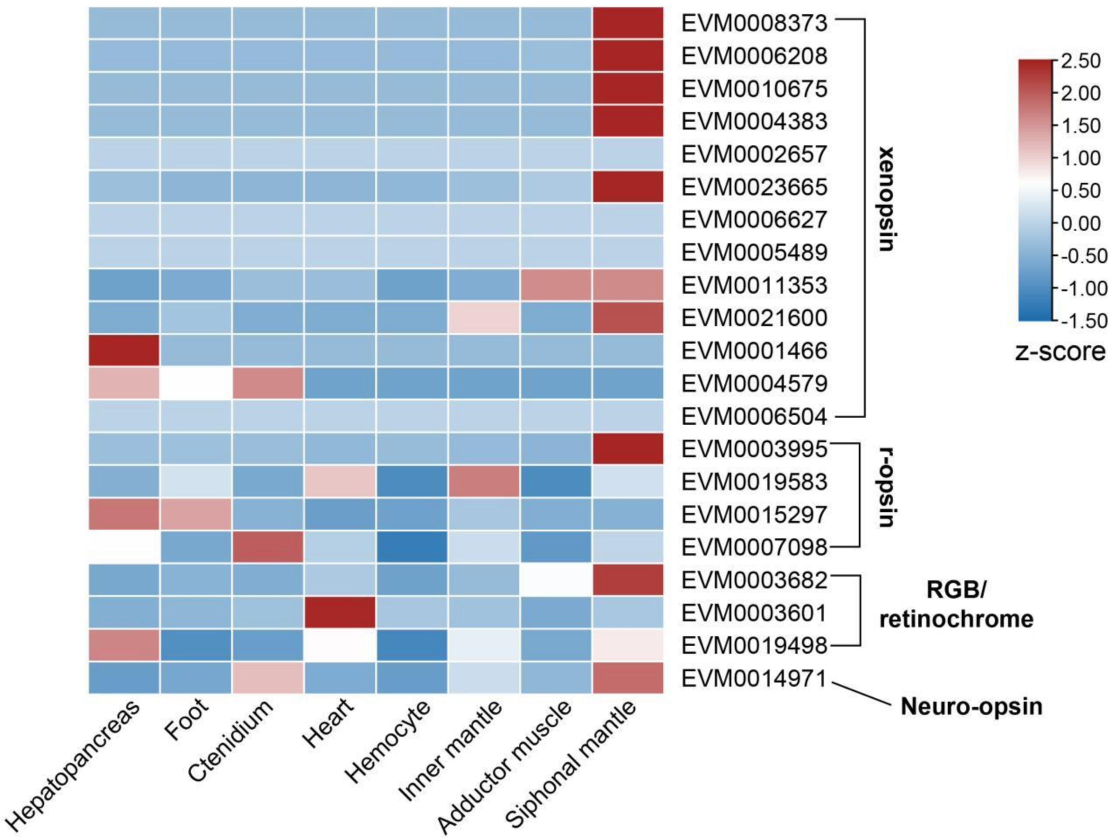


**Figure S22 Tissue distribution of opsin geneds in *T. squamosa*.** Heatmap shows the expression profile of opsin genes in different tissues. *x*-axis displays different tissues and *y*-axis shows the degree of expression of different opsin genes. Colored bars represent *z*-score calculated from RPKM-values of a target gene in different tissues.

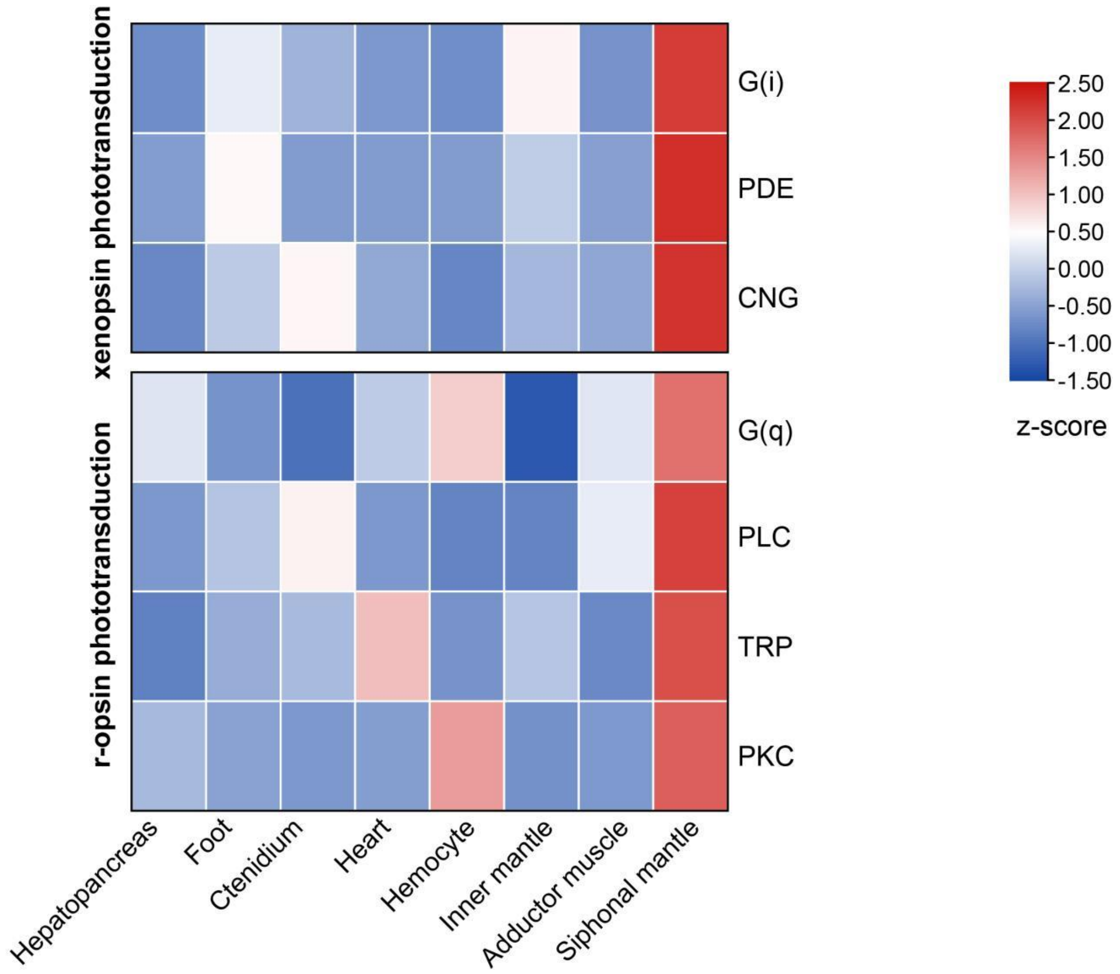


**Figure S23 Tissue distribution of gene sets for visual cycles and opsin signalling cascades.** Heatmap shows the expression profile of visual cycle genes in different tissues. *x*-axis displays different tissues and *y*-axis shows the degree of expression of different opsin genes. Colored bars represent *z*-score calculated from RPKM-values of a target gene in different tissues.

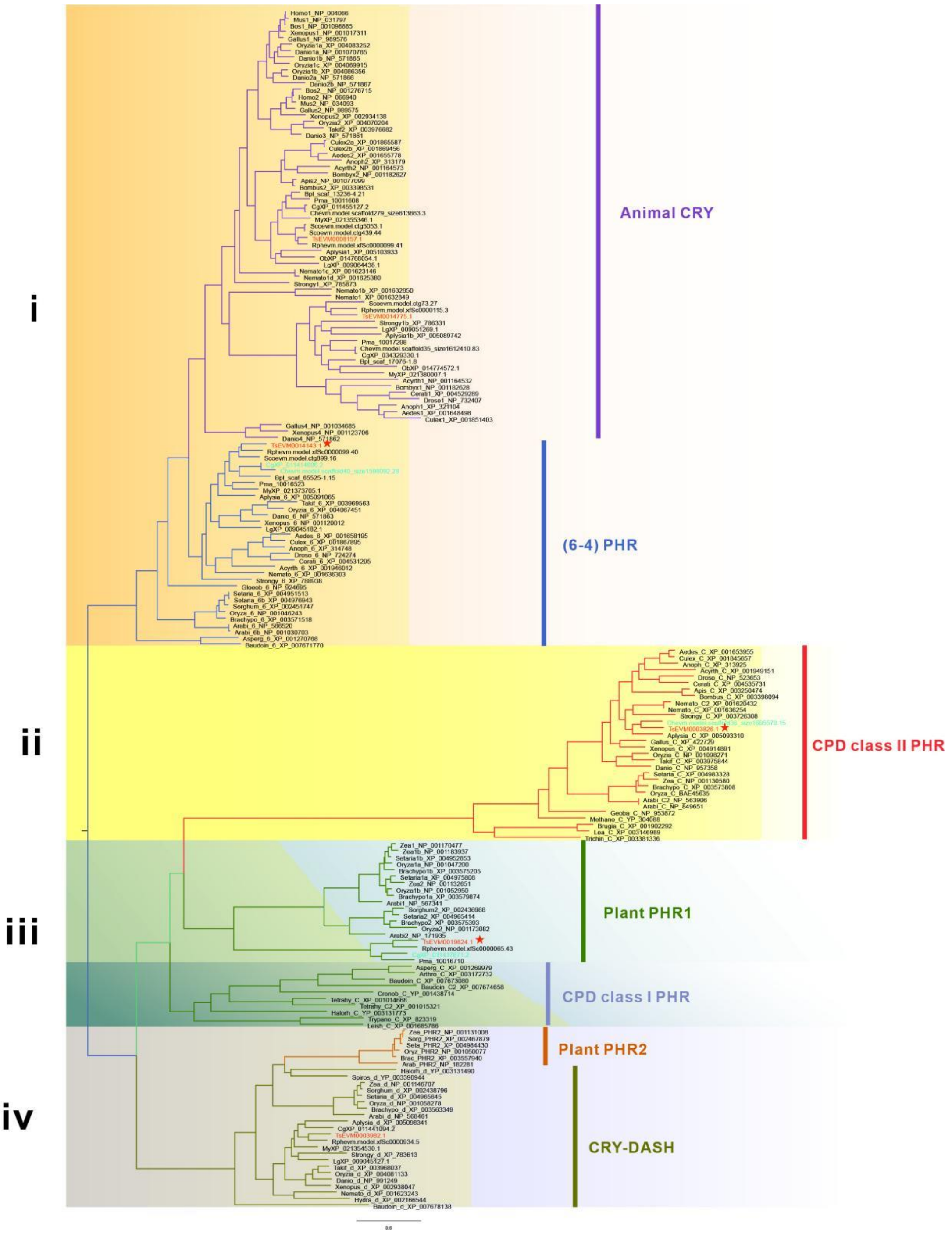


**Figure S24 Phylogenetic tree of the photolyase/cryptochrome family (CPF family).** Phylogenetic tree of CPF family is constructed by FastTree and visualized in FigTree. The background color indicates that these sequences belong to different subfamilies. Sequences of *T. squamosa* are displayed as red, and pentagram marked the genes significantly over-expressed in outer mantle. The leaves colored blue are photolyase orthologs in proximate bivalves (non-symbiotic bivalve *Crassostrea* oysters)

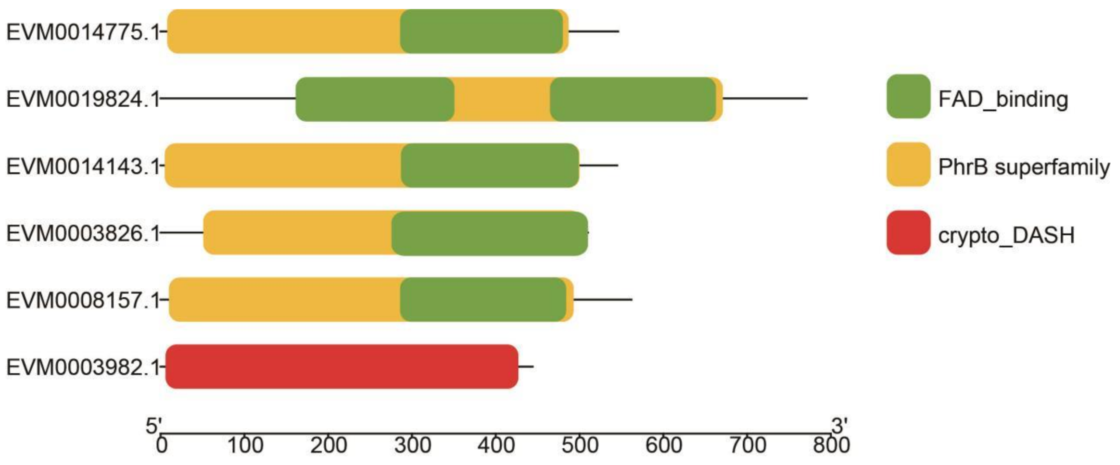


**Figure S25 Domain composition of CPF family members in *T. squamosa*.** Domain architecture was predicted by NCBI Conserved Domain Search and visualized by TBtools software.

A

| | Tridacna squamosa | Crassostrea gigas | Crassostrea hongkongensis |
|---|---|---|---|
| PHR1 | *Ts*EVM0014143 | *Cg*XP_011417671.2 | *Ch*evm.model.scaffold40_size1598092.28 |
| PHR3 | *Ts*EVM0019824 | *Cg*XP_011414696.2 | - |
| PHR2 | *Ts*EVM0003826 | - | *Ch*evm.model.scaffold36_size1605579.15 |

B

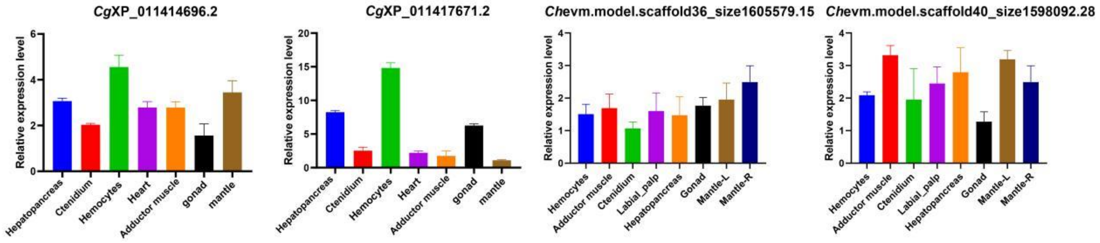


**Figure S26 Analysis of photolyase orthologs in proximate bivalves (non-symbiotic bivalve Crassostrea oysters)**. A. The table shows the photolyase orthologs correspondence based on the evolutionary tree between *T. squasoma*, *Crassostrea hongkongenesis* and *Crassostrea gigas*. B. Expression level of photolyase orthologs in oyster are examined by real-time quantitative PCR in different tissues, while no specific expression pattern could be observed.

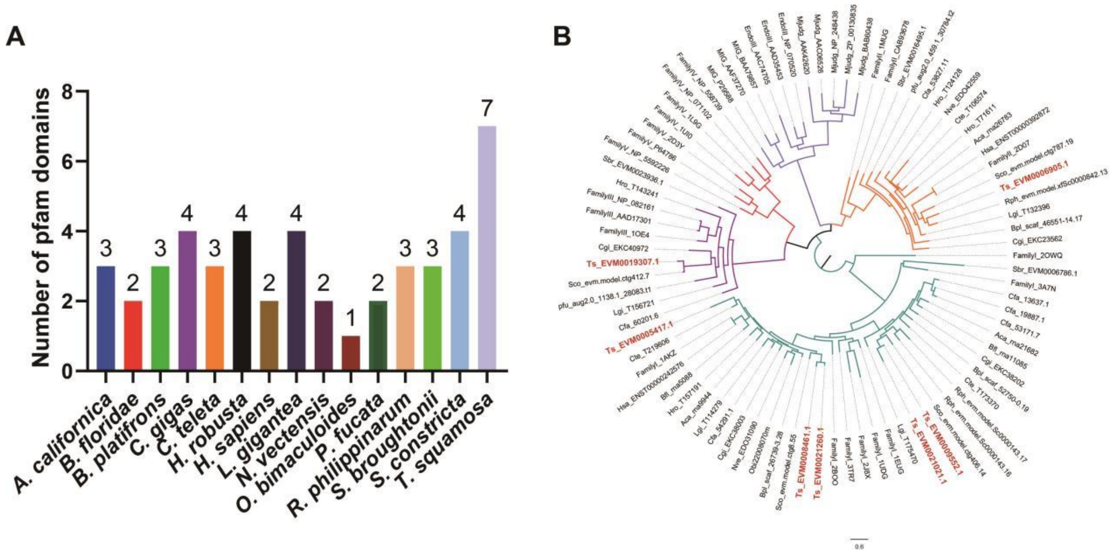


**Figure S27 Analysis of uracil-DNA glycosylase (UDG).** A. Histograms of the UDG gene numbers in analyzed species. The gene number is determined by the number of pfam domain. B. Evolutionary tree of UDG family among different species is constructed by FastTree and visualized in FigTree. Sequences identified in *T. squamosa* indicated in red color.

1. Simakov O, Marletaz F, Yue JX, O'Connell B, Jenkins J, Brandt A*, et al.* Deeply conserved synteny resolves early events in vertebrate evolution. *Nature ecology & evolution* 2020, **4**(6)**:** 820-830.
